# Denatured Albumin Gains a Function of Regulating Platelet Activity

**DOI:** 10.64898/2026.03.30.715112

**Authors:** Vivek Pandey, Subhankar Kundu, Sarah J Wang, Arghajit Pyne, Long Que, Xuefeng Wang

**Affiliations:** Hoxworth Blood Center, College of Medicine, University of Cincinnati, Cincinnati, OH, USA; Department of Electrical and Computer Engineering, Iowa State University, Ames, IA, USA; The department of Pathology and Laboratory Medicine, College of Medicine, University of Cincinnati, Cincinnati, OH, USA

## Abstract

Platelets are blood cells that play a critical role in hemostasis and thrombosis. Serum albumin, which constitutes approximately 50% of plasma proteins, has traditionally been considered non-interactive with platelets and not involved in platelet function. Here, using a molecular force sensor and platelet adhesion and aggregation assays, we show that serum albumin, if denatured, specifically binds to integrin α_IIb_β_3_ in platelets and transmits platelet contractile forces, regulating platelet activation, adhesion and aggregation as effectively as fibrinogen (the clotting factor I). 0.1% ethanol or 10 µM hydrogen peroxide, physiologically attainable in blood through alcohol intake or disease-elevated oxidative stress, are sufficient to denature albumin into a functional platelet ligand. These findings revealed albumin as a previously unrecognized but important regulator of platelet functions, with broad implications for the thrombotic risk in the context of physiological conditions that induce albumin denaturation.

## Introduction

Blood plasma contains hundreds of different proteins, with albumin and fibrinogen accounting for approximately 50% ^1^ and 5% ^2^ of total plasma proteins in healthy human adults, respectively. As the major coagulant protein, fibrinogen and other clotting factors interact with platelets and play a crucial role in hemostasis and thrombosis ^3^. By binding to the activated integrin α_IIb_β_3_ (glycoprotein IIb/IIIa) on platelets ^4–6^, fibrinogen mediates platelet aggregation at sites of vascular injury or thrombosis during primary hemostasis ^7^, and subsequently forms the fibrin meshwork of the blood clot during secondary hemostasis ^8^.

While the importance of fibrinogen in hemostasis and thrombosis is well established, serum albumin, the most abundant protein in blood plasma, is generally considered inert in platelet function, aside from its recognized roles in maintaining colloidal osmotic pressure ^9^ and transporting various substances ^10^ in the blood. During *in vitro* platelet assays, albumin has been frequently used as a surface blocking agent to reduce platelet activation and adhesion ^11–14^, reflecting its presumed inertness. Although several studies reported platelet adhesion on surface-adsorbed albumin ^15–17^, and attributed this effect to potential albumin structural alteration that leads to albumin-platelet interaction, these studies did not thoroughly exclude the possibilities that platelet adhesion arises from trace contaminants, such as fibrinogen or fibronectin, or surface electrochemical property altered by adsorbed proteins. Furthermore, platelets themselves can secrete fibrinogen, which may deposit on surfaces ^18, 19^ and mediate platelet adhesion. Overall, the mainstream view holds that albumin does not interact with platelets or directly influence hemostasis or thrombosis. The concept of a direct albumin-platelet interaction remains disregarded by the research field.

In this study, we investigated the interaction between albumin and platelets using a repurposed fluorescence-based force sensor to probe albumin-platelet binding with high sensitivity and specificity. Our experiments showed that denatured albumin, but not native one, binds specifically to integrin α_IIb_β_3_ on platelets and promotes platelet adhesion and aggregation as effectively as fibrinogen. Importantly, albumin denaturation can occur at the soluble state under physiological conditions, including exposure to low levels of ethanol or hydrogen peroxide. These findings identify a previously unrecognized role of albumin in platelet function and suggest a mechanism linking albumin denaturation to thrombosis under conditions such as alcohol exposure and disease-derived oxidative stress.

## Results

### Surface-adsorbed albumin supports platelet activation and adhesion

Previous studies have reported conflicting results on whether surface-adsorbed albumin supports platelet adhesion. To resolve this discrepancy, we tested platelet adhesion on albumin-or casein-coated glass surfaces and compared the result to HeLa cell adhesion under the same conditions.

To minimize the impact of sample variation to platelet adhesion test, one glass surface was coated with humans serum albumin by physical adsorption in a region, and the adjacent region was coated with casein, another commonly used blocking protein ^20^. Albumin was fluorescently labeled to distinguish the two regions by fluorescence imaging. Platelets activated with 10 µM ADP (adenosine diphosphate) and suspended in serum-free medium were applied to the entire surface. Following a 30 min incubation, F-actin imaging shows that platelets adhered and spread exclusively on albumin-adsorbed region, whereas no adhesion was observed on casein-adsorbed region (Fig. 1A). Similar results were obtained when albumin and casein were micropatterned on the same glass surface (Supplementary Fig. 1), confirming that platelet adhesion was specific to albumin-adsorbed regions.

**Fig. 1.**
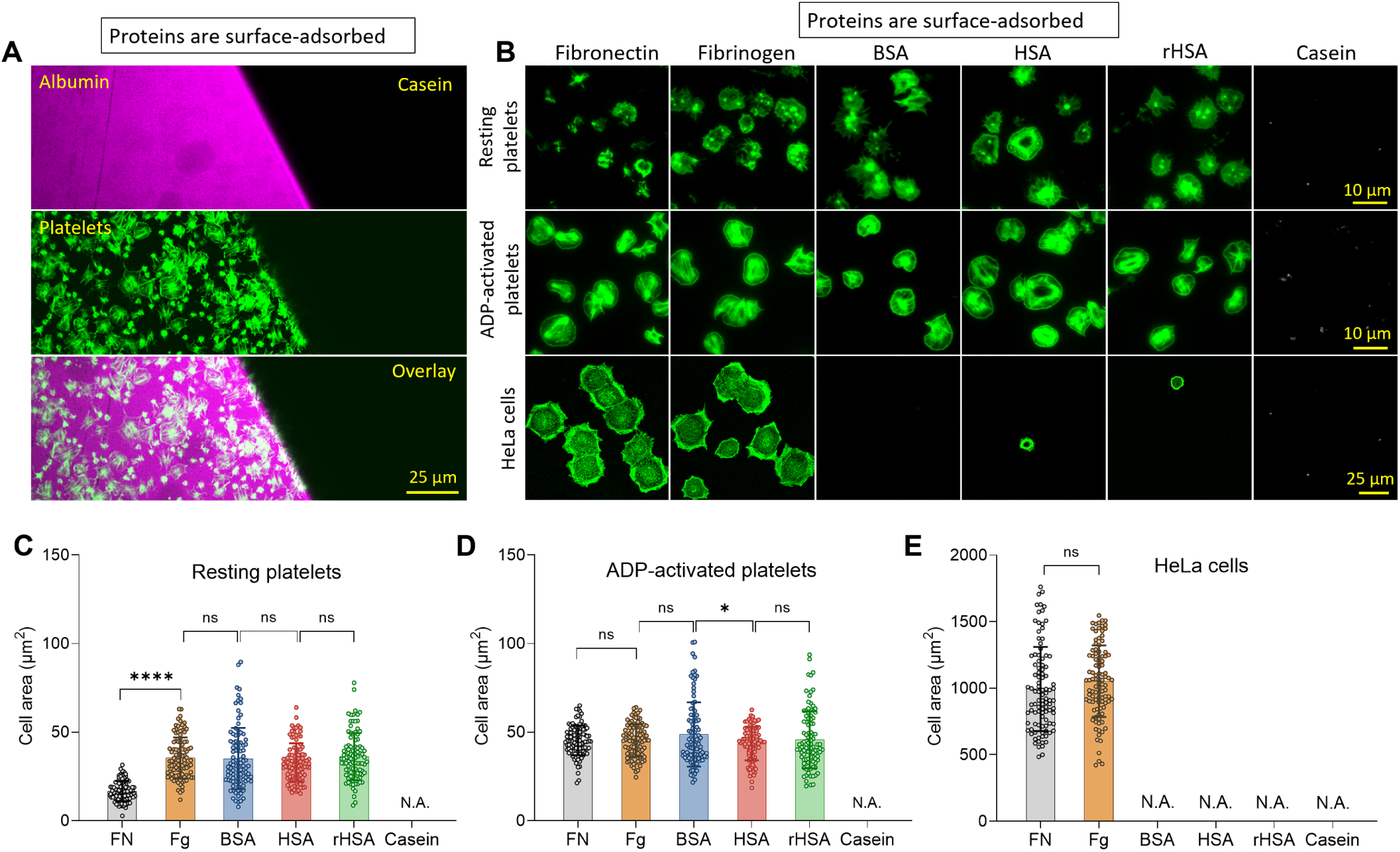
Surface-adsorbed albumin supports platelet activation and adhesion. (A) Platelets (green) adhered exclusively to the albumin-adsorbed region (magenta), but not to the casein-adsorbed region (dark). Albumin was pre-labeled with Alexa488. Platelets were stained with phalloidin to label F-actin. (B) Resting platelets, ADP-activated platelets and HeLa cells on fibronectin (FN), fibrinogen (Fg), bovine serum albumin (BSA), human serum albumin (HSA), recombinant human serum albumin (rHSA) and casein-adsorbed glass surfaces, respectively. (**C-E**) Cell spreading areas of platelets or HeLa cells on the six surfaces. Platelets in (C) were kept at a resting state prior to cell plating, whereas platelets in (D) were pre-activated by 10 µM ADP. (**** *P*<0.0001; ns *P*>0.05; Each data point represents one cell; For each condition, N = 100 cells from three independent experiments; Error bars indicate SD).

Because albumin is generally purified from plasma, it may contain trace amounts of fibrinogen and fibronectin, both of which promote platelet adhesion when surface-bound ^21, 22^. We therefore considered the possibility that residual levels of these proteins in albumin products may account for the observed platelet adhesion. To test this, we examined HeLa cell adhesion on albumin-adsorbed surfaces. The rationale was that if fibrinogen or fibronectin were present in the adsorbed albumin, the surface should also support HeLa cell adhesion. We also tested recombinant human serum albumin (rHSA) which is not expected to contain fibrinogen or fibronectin. Glass substrates were adsorbed with ultrapure bovine serum albumin (BSA), ultrapure human serum albumin (HSA), and rHSA, respectively. In addition, fibronectin-and fibrinogen-adsorbed surfaces were included as positive controls, while casein-adsorbed surfaces served as a negative control. Resting platelets, ADP-activated platelets and HeLa cells were incubated on these surfaces. As shown in Fig. 1B, resting platelets became activated and adhered to BSA-, HSA-, rHSA-, fibrinogen-, and fibronectin-adsorbed surfaces, but not to casein-adsorbed surfaces. ADP-activated platelets exhibited similar adhesion outcomes to resting platelets on these surfaces. In contrast, HeLa cells adhered and spread robustly on fibrinogen-and fibronectin-coated surfaces but had no adhesion on BSA-, HSA-, rHSA-or casein-coated surfaces. This indicates that the albumin products used in this work do not contain fibrinogen or fibronectin at levels sufficient to support cell adhesion, particularly in the case of rHSA, which was derived from cultured cells rather than plasma. Quantitative analysis (Figs. 1C-1E) showed that resting platelets spread to an average area of ∼35 µm² on fibrinogen, BSA, HSA, rHSA, but exhibited reduced spreading on fibronectin (∼50% of the area on the other substrates). ADP-activated platelets spread to ∼45 µm² on all five substrates without significant differences. HeLa cells spread to ∼1000 µm² on fibronectin-or fibrinogen-coated surfaces. Cell area quantification algorithm is described in Supplementary Fig. 2. These results confirm that surface-adsorbed albumin indeed supports platelet activation and adhesion, suggesting that surface adsorption may induce albumin structural change, or cause albumin denaturation that exposes platelet-binding sites.

### Denatured albumin supports platelet adhesion

Above tests still cannot rule out other possible mechanism, e.g., that albumin modifies the physicochemical properties of the glass, such as surface charge or hydrophobicity, thereby rendering it more conducive to platelet adhesion. To verify that denaturation is the key and sufficient process enabling albumin-platelet interaction, we tethered native albumin (Alb) or heat-denatured albumin (dnAlb) via biotin-avidin coupling on PEGylated non-fouling surfaces, which are functionalized with polyethylene glycol (PEG) polymer that forms molecular brush-like structure. This PEG structure suppresses physical adsorption of proteins on the surface and retains protein original structures ^23^.

As shown in Fig. 2A, biotin-labeled human serum albumin (Alb-biotin), denatured albumin (dnAlb-biotin), fibrinogen (Fg-biotin), and the RGD (Arg-Gly-Asp) peptide (RGD-biotin) were tethered to PEGylated glass surfaces. dnAlb-biotin was prepared by heating Alb-biotin solution at 90°C for 10 min. RGD is the critical integrin-binding peptide motif in fibrinogen and fibronectin ^24^. Platelets were plated on these surfaces and incubated for 30 min. F-actin imaging revealed poor platelet adhesion on Alb-tethered surfaces, but robust adhesion (platelet spread) on dnAlb-, Fg-, and RGD-tethered surfaces (Fig. 2B), indicating that denaturation converts albumin into a platelet ligand that, if surface-tethered, supports platelet adhesion. In Supplementary Fig. 3, we further demonstrated that dnAlb has minimal or no binding capacity with fibrinogen, ruling out the possibility that dnAlb binds platelet-released fibrinogen to indirectly support platelet adhesion.

**Fig. 2.**
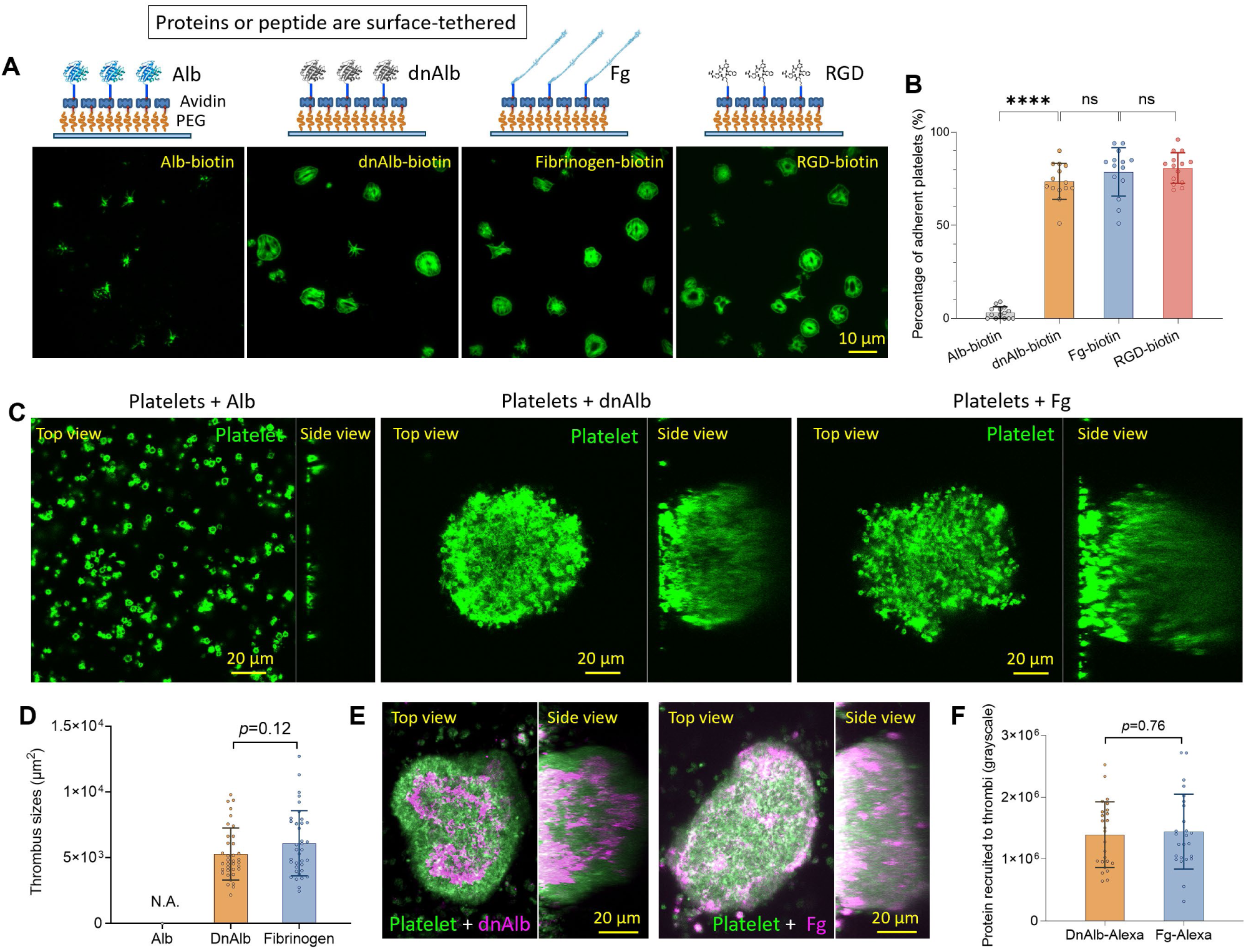
Denatured albumin supports platelet adhesion and aggregation. (A) Four biotin-labeled molecules: albumin (Alb), heat-denatured albumin (dnAlb), fibrinogen, and RGD peptide, were tethered to non-fouling PEGylated glass surfaces via biotin-avidin interaction. ADP-activated platelets were incubated on these surfaces for 30 min. (B) Percentages of platelets that spread on the four surfaces. (Each data point was calculated over one imaging field; N = 10 images for each condition.) (C) Platelet aggregates (thrombi) were formed in platelet solutions supplemented with 50 µg/mL dnAlb or fibrinogen, but not in 50 µg/mL Alb-supplemented one. Thrombi was reported by F-actin staining and imaging. (D) The sizes of thrombi formed in (H) (For each condition, N = 36 thrombi from three independent experiments). (E) Co-imaging of dnAlb and platelets, or Fg and platelets, in thrombi. The dnAlb and Fg were labeled with Alexa488. (F) Fluorescent intensities of dnAlb and fibrinogen recruited to thrombi. The dnAlb and fibrinogen were labeled with Alexa488. (N = 25 thrombi for each condition)

### Denatured albumin mediates platelet aggregation as effectively as fibrinogen

As fibrinogen in the soluble state is the key protein mediating platelet aggregation ^25^, we next tested whether soluble dnAlb (denatured by heating at 90 °C for 10 min) can induce platelet aggregation similarly to fibrinogen. Purified platelets were resuspended in serum-free medium and supplemented with 50 µg/mL native human serum albumin (Alb), dnAlb, or fibrinogen, followed by sample shaking (emulating fluidic stress) at 37 °C for 30 min. The platelet samples were then transferred to Petri dishes to examine thrombus formation. As shown in Fig. 2C and Supplementary Fig. 4, Alb supplementation did not induce thrombus formation, whereas dnAlb and fibrinogen both promoted the formation of thrombi with comparable sizes (Fig. 2D). To verify dnAlb incorporation into thrombi, in another test, all platelet suspensions were supplemented with dye-labeled dnAlb or fibrinogen, respectively. Supplementary Movie 1 shows that dnAlb was indeed recruited to thrombi. Fluorescence imaging (Fig. 2E and Supplementary Fig. 5) and quantification (Fig. 2F) demonstrated recruitment of dnAlb and fibrinogen into thrombi at similar levels. Together, these results confirm that dnAlb mediates platelet aggregation as effectively as fibrinogen.

### Denatured albumin transmits platelet contractile forces

To directly investigate albumin-platelet interactions, we developed an albumin-conjugated integrative tension sensor (ITS). ITS is a surface-tethered molecular force probe that can be conjugated with specific ligands to visualize membrane receptor-transmitted cellular forces ^26–28^. In previous work, we used RGD-conjugated ITS to examine platelet force generation via integrin α_IIb_β_3_ ^28, 29^. Here, we replaced the RGD ligand with albumin to specifically assess platelet-albumin interactions.

As shown in Fig. 3A, an ITS construct consists of an 18-base-paired double-stranded DNA (dsDNA). The upper DNA strand was conjugated with a black hole quencher (BHQ2) and a ligand molecule, here a human serum albumin, via copper-free click chemistry. The conjugation was performed in a physiologically compatible buffer to preserve albumin’s structural integrity (see Supplementary Fig. 6 for details). It was also ensured that each albumin molecule was conjugated with no more than one DNA strand (Supplementary Fig. 7). The lower DNA strand was labeled with an Atto647N fluorophore and a biotin tag, enabling immobilization of ITS on avidin-presenting surfaces. During experiments, the dsDNA-based ITS is densely tethered on surfaces at >1000 units/µm². Cells adhere to the surface and engage the ligand presented on the ITS. When the receptor-ligand bond transmits force that exceeds a critical threshold ^30^, the dsDNA dissociates, freeing the fluorophore from quenching and thereby providing direct fluorescent readout of a force event. Albumin-conjugated ITS (Alb-ITS) is thus capable of probing platelet-albumin interactions by fluorescence imaging with high sensitivity and specificity.

**Fig. 3.**
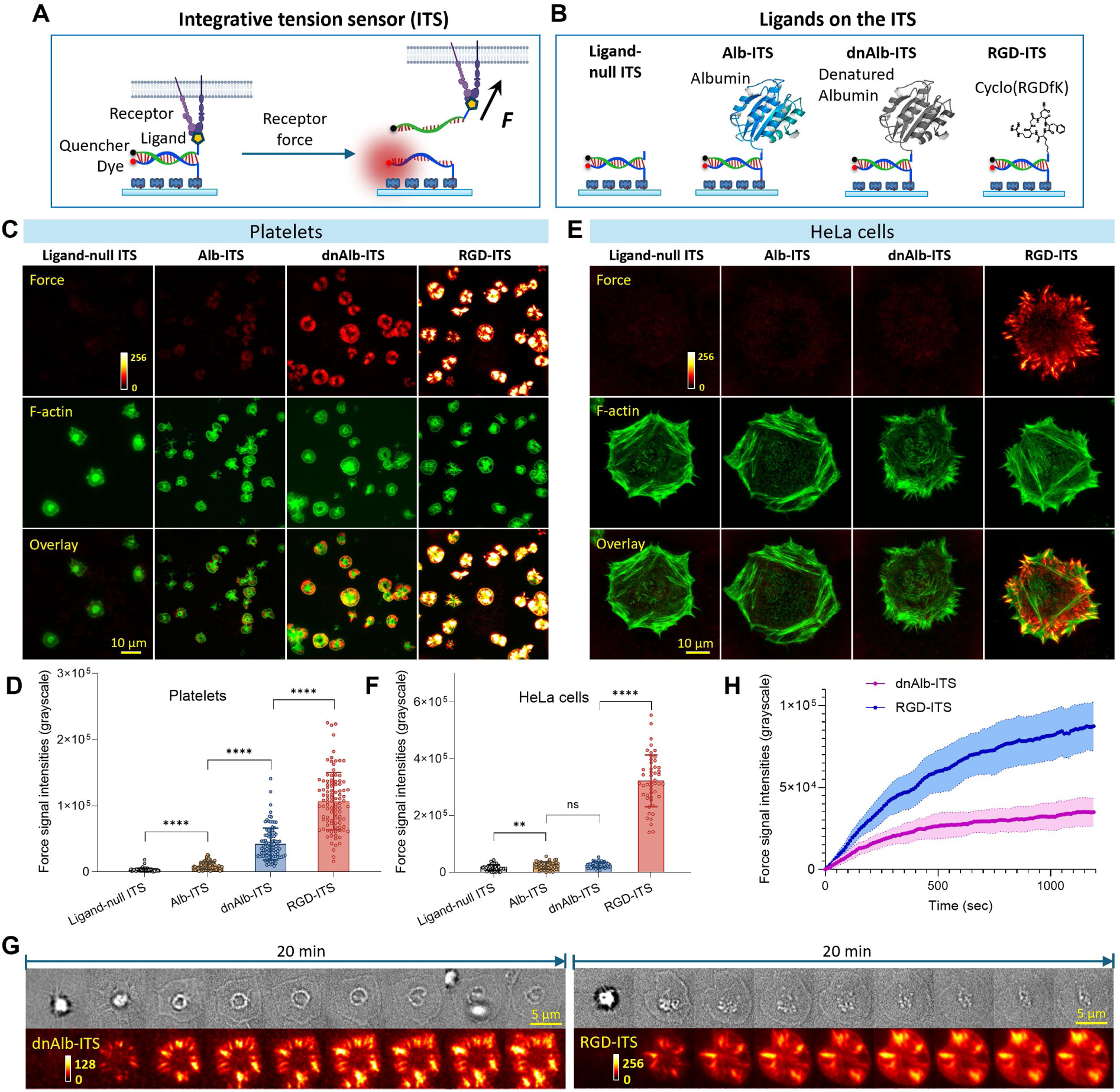
Denatured albumin transmits platelet-generated forces during platelet adhesion. (A) Schematics and working principle of integrative tension sensor (ITS). (B) Schematics of ligand-null ITS and ITS constructs equipped with albumin (Alb-ITS), denatured albumin (dnAlb-ITS, heated at 90°C for 10 min) and integrin peptide ligand RGD (RGD-ITS). (C) Platelets on the surfaces grafted with ligand-null ITS, Alb-ITS, dnAlb-ITS and RGD-ITS, respectively. F-actin and platelet adhesive force reported by ITS were imaged and displayed. (D) Platelet force intensity analysis (Each data point represents one platelet; N = 100 platelets from three independent experiments). (E) HeLa cells incubated on the four ITS surfaces. (F) HeLa cell force intensity analysis (\*\*\*\**P*<0.0001; \*\**P*<0.01; ns; *P*>0.05; Each data point represents one cell; N = 50 cells from three independent experiments). (G) Live-cell force imaging over individual platelets plated on a dnAlb-ITS surface or an RGD-ITS surface, respectively. Related to Supplementary Movies 2 and 3. (H) Force signal intensities in single platelets accumulated over time on the dnAlb-ITS and RGD-ITS surfaces, respectively. Each curve represents the mean force signal from 9 platelets, with the shading denoting SD.

In experiments, four surfaces were immobilized with ligand-null ITS (negative control), Alb-ITS, dnAlb-ITS (denatured by heating) and RGD-ITS (positive control), respectively (Fig. 3B). ADP-activated platelets were incubated on these four surfaces for 30 min. The samples were then washed, fixed and stained for the co-imaging of platelet forces and F-actin structures. As shown in Fig. 3C, negligible force signals were observed on the ligand-null ITS surface, low-level force signals were observed on the Alb-ITS surface, substantially higher force signals were observed on the dnAlb-ITS surface, and the highest force signals were observed on the RGD-ITS surface. Data analysis in Fig. 3D shows that platelet force signals on the dnAlb-ITS surface are significantly higher than those on the Alb-ITS surface, and on average approximately half of those observed on the RGD-ITS surface. This experiment demonstrates that dnAlb directly interacts with platelets and transmit platelet forces in a manner similar to integrin ligand RGD peptide. In contrast, HeLa cells did not produce force signals on ligand-null ITS, Alb-ITS or dnAlb-ITS surfaces, but produced force signals normally on the RGD-ITS surface (Figs. 3E and 3F). This comparison suggests that dnAlb-cell interaction is specific for platelets.

Live-cell imaging were performed on platelets on dnAlb-ITS and RGD-ITS surfaces, respectively (Fig. 3G, Supplementary Movies 2 and 3). The results show similar platelet force patterns on these two surfaces, exhibiting streak and plaque-like features ^29^. The majority of force signals were accumulated in the first 10 min after platelet adhesion (Fig. 3H). The distinct spatial pattern of platelet forces on the dnAlb-ITS surface indicates that the dnAlb-platelet interaction is a regulated process rather than nonspecific binding.

### Denatured albumin specifically binds to integrin α_IIb_β_3_ in platelets

We next investigated potential membrane receptors and adaptor proteins involved in dnAlb-transmitted platelet forces. Platelets adhering to dnAlb-ITS surfaces were immunostained with antibodies against integrin β_3_, integrin β_1_, vinculin, and phosphorylated myosin light chain (pMLC). As shown in Fig. 4A, F-actin, target proteins and dnAlb-transmitted forces were co-imaged. Line profile analysis (Fig. 4B) was performed to evaluate spatial co-localization between the target proteins and force signals. Both the images and line analysis demonstrated strong co-localization of dnAlb-transmitted forces with integrin β_3_ and vinculin, suggesting that dnAlb likely engages integrin α_IIb_β_3_ and enables platelet force transmission.

**Fig. 4.**
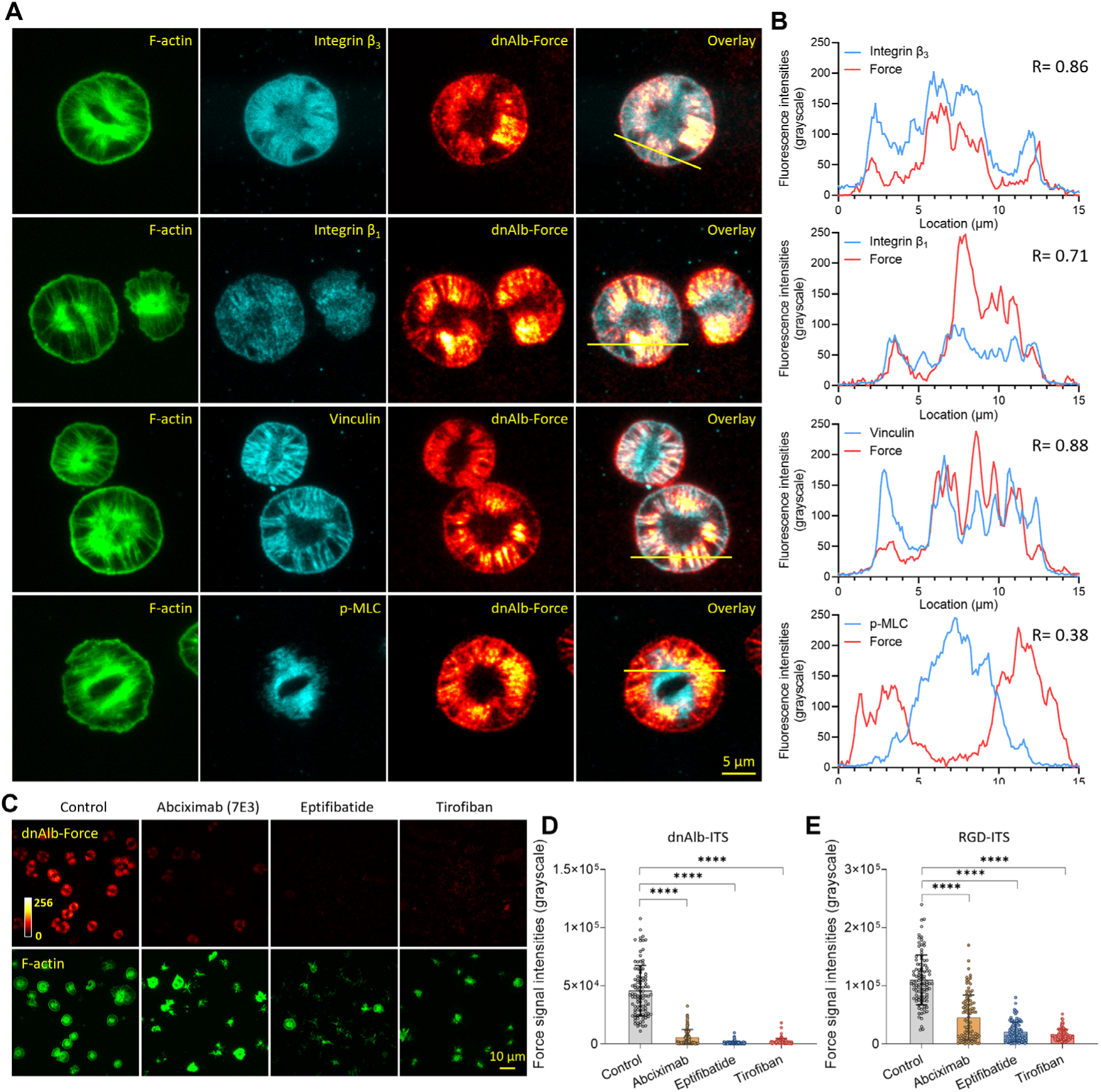
Denatured albumin specifically binds to integrin α_IIb_β_3_ in platelets during force transmission. (A) Co-imaging of F-actin, integrin β_3_, integrin β_1_, vinculin, p-MLC and dnAlb-transmitted platelet forces on dnAlb-ITS surfaces. (B) Line profiles of protein signal and force signal. The line profiles are marked yellow in (A). Cross-correlation values of proteins and force signals are indicated in the graphs. (C) F-actin and dnAlb-transmitted platelet forces in platelets treated with three inhibitors of integrin α_IIb_β_3_. (**D**-**E**) Quantifications of dnAlb-transmitted platelet forces and RGD-transmitted platelet forces with integrin α_IIb_β_3_ inhibited. (Each data point represents one cell; For each condition, N = 100 platelets from three independent experiments).

To confirm that dnAlb specifically interacts with integrin α_IIb_β_3_ in platelets, we examined the effect of integrin α_IIb_β_3_ inhibition on platelet forces on dnAlb-ITS surfaces. Three inhibitors of integrin α_IIb_β_3_ were tested: tirofiban, eptifibatide, and abciximab. Tirofiban ^31^ is a well-characterized small-molecule antagonist, while eptifibatide ^32^ is a cyclic heptapeptide derived from snake venom that mimics the RGD motif. Both agents bind to the ligand-binding pocket of integrin α_IIb_β_3_, thereby preventing ligand engagement. Abciximab ^33^, in contrast, is a chimeric monoclonal antibody Fab fragment that binds with high affinity to integrin α_IIb_β_3_ and sterically blocks ligand binding. All three inhibitors are established antiplatelet therapies used clinically to reduce thrombotic cardiovascular events ^34, 35^. As shown in Fig. 4C and Supplementary Fig. 8A, platelets treated with tirofiban or eptifibatide were practically unable to produce force signals on dnAlb-ITS surfaces, and platelets treated with Abciximab produced significantly lower force signals than platelets of control samples (Fig. 4D). The efficacies of these inhibitors over platelet force suppression were also verified on RGD-ITS surfaces (Fig. 4E and Supplementary Fig. 8B).

Collectively, these experiments prove that denatured albumin specifically binds to integrin α_IIb_β_3_ to mediate platelet adhesion and force transmission.

### Source and range of dnAlb-transmitted molecular forces in platelets

Using dnAlb-ITS, we calibrated the source and magnitude of platelet molecular forces transmitted through dnAlb. Because myosin II is a primary driver of platelet adhesive forces ^36^, we assessed its contribution by inhibiting myosin II activity. As shown in Figs. 5A and 5B, platelets treated with 25 µM blebbistatin remained adhered and spread but exhibited significantly reduced force signals on the dnAlb-ITS surface compared with controls, confirming myosin II as the primary source of dnAlb-transmitted platelet forces.

**Fig. 5.**
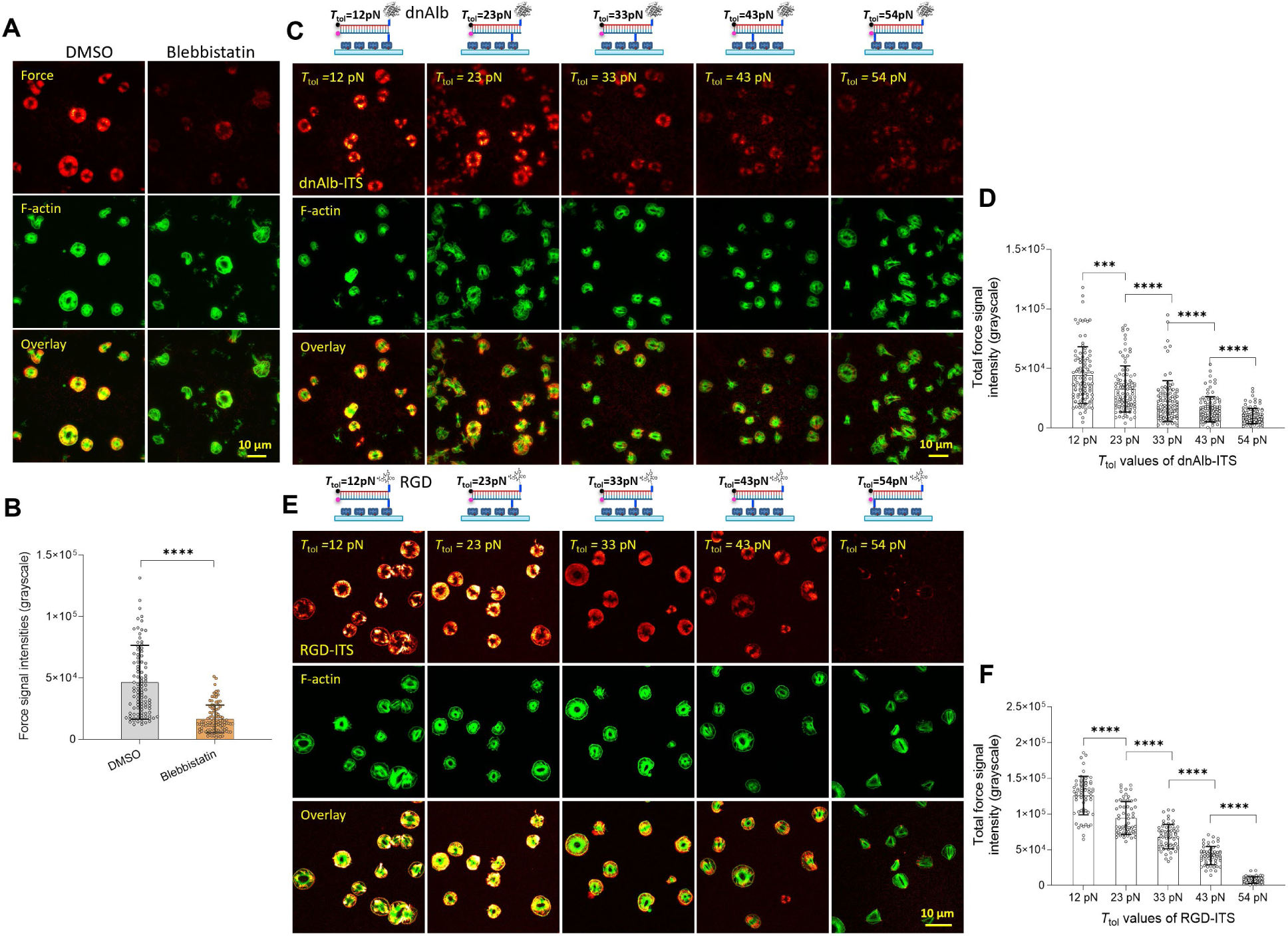
Source and magnitude range of dnAlb-transmitted molecular force in platelets. (A) dnAlb-transmitted platelet forces with platelets treated with DMSO (vehicle control) or 25 µM blebbistatin. (B) Quantification of force signal intensities in (A). (\*\*\*\**P*<0.0001; Each data point represents one cell; For each condition, N = 100 platelets from three independent experiments) (C) Platelet forces imaged on dnAlb-ITS surfaces with tension tolerance (*T*_tol_) = 12, 23, 33, 43 or 54 pN, respectively. *T*_tol_ values indicate the force thresholds required to activate the ITS to the fluorescent state. (D) Quantification of force signal intensities in (C). (\*\*\*\**P*<0.0001; Each data point represents one cell; For each condition, N = 100 platelets from three independent experiments). (E) Platelet forces imaged on RGD-ITS surfaces with tension tolerance (*T*_tol_) = 12, 23, 33, 43 or 54 pN, respectively. (F) Quantification of force signal intensities in (E). (\*\*\*\**P*<0.0001; Each data point represents one cell; For each condition, N = 60 platelets from three independent experiments).

We also calibrated the molecular force level transmitted by dnAlb-integrin bonds during platelet adhesion. This is enabled by tuning the tension tolerance (*T*_tol_) of ITS constructs. *T*_tol_ indicates the critical force required to activate an ITS construct to fluorescent state, tunable in the range of 12 picoNetwen (pN) to 54 pN ^30, 37^. The *T*_tol_ value of ITS used in this study is 12 pN, except in this section. Here, a series of dnAlb-ITS and RGD-ITS surfaces were prepared, with *T*_tol_ = 12, 23, 33, 43 and 54 pN, respectively. As shown in Figs. 5C and 5D, platelet force signal intensities decrease monotonically with increasing *T*_tol_ values. On the 54 pN dnAlb-ITS surface, representing forces exceeding 54 pN, signal intensity is approximately 20% of that observed on the 12 pN dnAlb-ITS surface, suggesting that the magnitude of dnAlb-transmitted molecular forces in platelets is largely distributed within the 12-54 pN range. A similar trend was observed on RGD-ITS surfaces (Figs. 5E and 5F), indicating that the affinity between dnAlb and integrin α_IIb_β_3_ is comparable to that between RGD and integrin.

### Ethanol and H_2_O_2_ at physiologically levels can denature albumin into a platelet ligand

Having verified that heat-denatured albumin functions as an integrin ligand for platelets, we next examined whether ethanol and hydrogen peroxide, two physiologically relevant chemical factors, can similarly denature albumin into platelet-binding ligands. Ethanol is known to denature proteins by weakening hydrophobic interactions and disrupting hydrogen-bonding networks, leading to protein unfolding. Blood ethanol concentration during alcohol consumption can reach 0.4%, with a level around 0.4% considered potentially fatal ^38^. Reactive oxygen species (ROS), such as hydrogen peroxide (H_2_O_2_), is another potential factor causing protein denaturation ^39, 40^. In blood, H_2_O_2_ is typically present at 1-5 µM under normal conditions. Pathological or inflammation states, including diabetes, sepsis, neurodegeneration, and cancer, often exhibit elevated oxidative stress, with H_2_O_2_ levels reaching 10-50 µM or higher ^41^. With these considerations, we investigated whether ethanol and H_2_O_2_ at physiologically attainable levels could transform albumin to an integrin α_IIb_β_3_ ligand.

In the experiment, Alb-ITS was treated by heat, SDS (Sodium Dodecyl Sulfate), ethanol or H_2_O_2_, respectively (Fig. 6A and Supplementary Fig. 9). Compared to Alb-ITS surfaces, platelets generated clear force signals on all dnAlb-ITS surfaces. dnAlb-ITS prepared by heating at 60°C, 75°C, and 90°C for 10 min produced progressively stronger force signals (Fig. 6B). dnAlb-ITS heated at 90°C for 5 min, 15 min and 60 min yielded the strongest force signal with 15 min heating time (Fig. 6C and Supplementary Fig. 9A), suggesting that prolonged heating may over-denature albumin and reduce its affinity to integrin α_IIb_β_3_. 1% SDS-treated dnAlb-ITS also induced positive platelet signals, peaking at 15 min treatment (Fig. 6D and and Supplementary Fig. 9B).

**Fig. 6.**
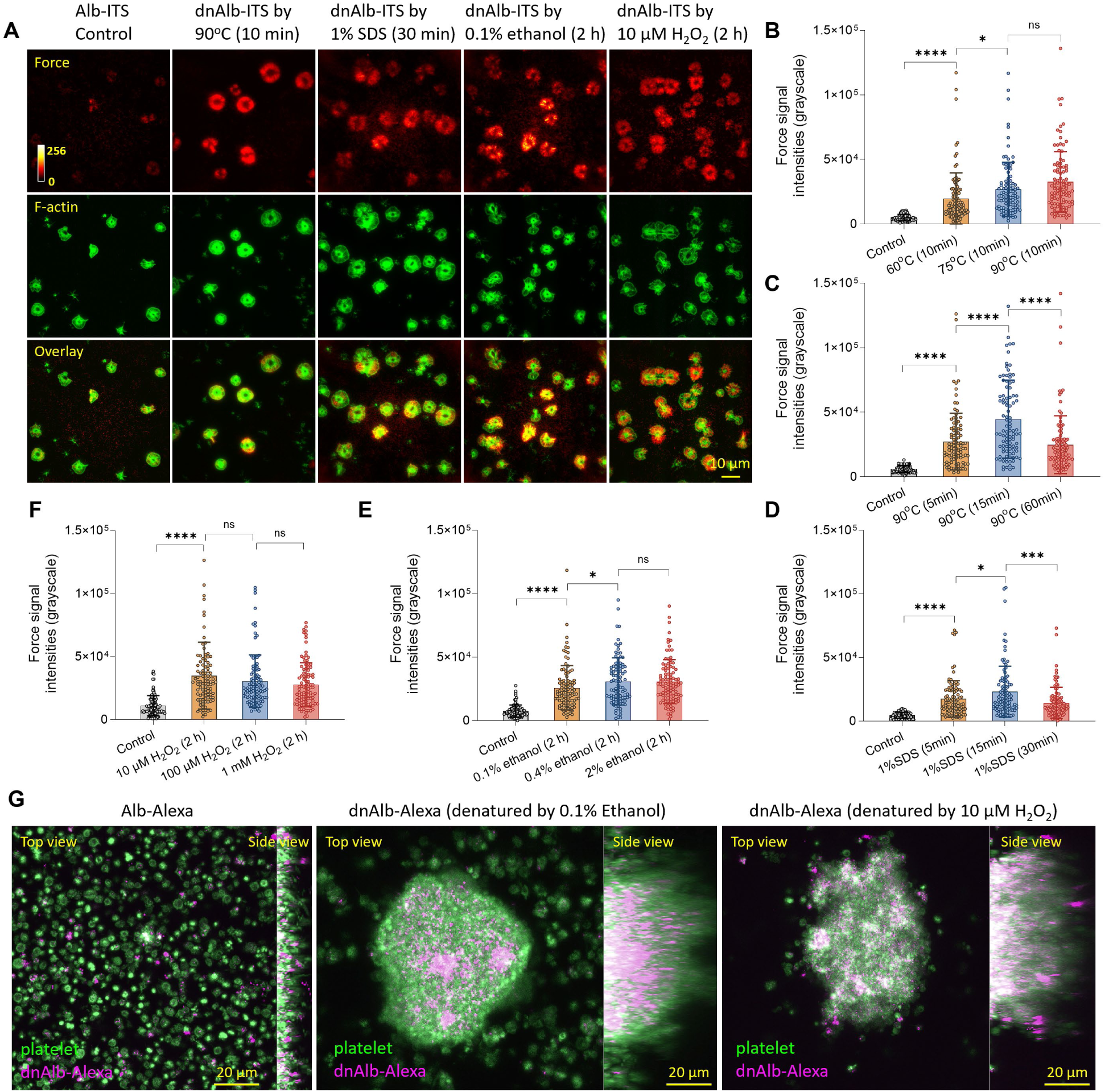
Albumin denatured by heating, SDS, ethanol or H_2_O_2_ becomes platelet ligands. (**A**) Force and F-actin of platelets plated on Alb-ITS (Control) and dnAlb-ITS surfaces. The dnAlb-ITS were either pre-denatured by heating at 90 °C for 10 min or treated with 1% SDS for 30 min, 0.1% ethanol for 2 h, or 10 µM H_2_O_2_ for 2 h, respectively. (**B-F**) Platelet force signal intensities on dnAlb-ITS surfaces. The Alb-ITS were pre-denatured by heating at different temperatures (B), heating at 90°C for different durations (C), treated with 1% SDS for different durations (D), treated with ethanol (E) or H_2_O_2_ (F) at different concentrations for 2 h. (For each condition, N = 100 platelets from three independent experiments). (G) Platelet aggregation in mixtures of platelets and Alexa488-labeled albumin (Alb-Alexa). Platelets showed no aggregation in native Alb-Alexa (50 µg/mL), but aggregated readily in Alb-Alexa pre-denatured with 0.1% ethanol or 10 µM H₂O₂. The dnAlb-Alexa (magenta) was also visible in the platelet aggregates (green).

Importantly, ethanol and H_2_O_2_ at physiologically tolerable levels similarly denatured albumin into an integrin α_IIb_β_3_ ligand. Platelets generated significant force signals on dnAlb-ITS pretreated with 0.1%, 0.4%, or 2% ethanol for 2 h (Fig. 6E and Supplementary Fig. 9C), as well as on dnAlb-ITS pretreated with 10 µM, 100 µM, or 1 mM H_2_O_2_ for 2 h (Fig. 6F and Supplementary Fig. 9D). These results indicate that albumin can be transformed into a functional platelet ligand under conditions relevant to human physiology.

We further observed that excessive denaturation ultimately abolished dnAlb’s ability to transmit platelet forces (Supplementary Fig. 10), indicating that the albumin-platelet interaction may not be driven by a specific peptide motif in albumin.

### Ethanol or H_2_O_2_-denatured albumin mediates platelet aggregation

Finally, we tested whether albumin denatured by ethanol or H_2_O_2_ at the low levels can mediate platelet aggregation. In the experiments, purified platelets resuspended in serum-free medium were supplemented with 50 µg/mL native Alexa488-labeled human serum albumin (Alb-Alexa), Alb-Alexa denatured by 0.1% ethanol or Alb-Alexa denatured by 10 µM H_2_O_2_, respectively. The platelet samples were shaken at 150 rpm at 37 °C for 30 min and then transferred to Petri dishes to examine thrombus formation and the recruitment of albumin to thrombi. As shown in Fig. 6G, native Alb did not induce thrombus formation, whereas both ethanol-denatured albumin and H_2_O_2_-denatured albumin induced the formation of thrombi. dnAlb, either induced by ethanol or H_2_O_2_, was shown to be incorporated into thrombi. These results confirm that human serum albumin, when exposed to ethanol or H_2_O_2_, even at physiologically tolerable levels, can be denatured into a platelet ligand capable of mediating platelet aggregation.

## DISCUSSION

Serum albumin, the most abundant protein in plasma, is traditionally considered biologically inert in the context of platelet function and associated hemostasis or thrombosis. In this study, we show that denatured albumin acts as a functional platelet ligand, specifically binding to integrin α_IIb_β_3_ in platelets and supporting platelet activation, adhesion and aggregation as effectively as fibrinogen.

It is not rare that denaturation can help a protein gain new function. The prion protein, if misfolded, can catalyze new prion protein to misfolded state and amplify the protein denaturation ^42^. Heat-shock protein Hsp33, under oxidative stress, exposes hydrophobic patches that bind and protect unfolded client proteins ^43^. As a globular protein with high conformational flexibility, albumin is sensitive to environmental condition and prone to undergo conformational changes ^44^. The high affinity and specificity of denatured albumin for integrin α_IIb_β_3_ in platelets, inducible by physiologically relevant factors, suggest that albumin can acquire a functional role in platelet activity through denaturation. As platelets are constantly exposed to albumin, this function may represent an evolutionarily conserved adaptation rather than a random occurrence.

While the exact structural basis and *in vivo* process of dnAlb-platelet interaction remain to be elucidated, this finding already holds significant implications for our understanding of platelet biology and physiology. Given the high abundance of albumin in blood and its readily conversion into a platelet ligand under physiologically relevant levels of alcohol or H_2_O_2_, denatured albumin likely plays a significant role in hemostasis and thrombosis. Our work offers a new perspective on thrombosis etiology, as many conditions may denature albumin and thereby impact platelet function. Previous studies indicate that albumin denaturation can occur in the bodies, for instances, due to oxidative stress ^45^, liver diseases ^46^, aging ^47^, etc. In addition, factors such as fever ^48^, abnormal pH ^49^, heavy metal exposure ^50^ may also potentially denature albumin. Albumin denaturation may therefore represent a previously overlooked factor linking diverse pathological conditions to heightened thrombotic risk.

## Methods

### Ethical statement

The use of human platelet samples in this study was reviewed and approved by the Institutional Biosafety Committee (IBC) at the University of Cincinnati under the protocol IBC#: 23-08-04-23. These platelet samples were categorized as “human specimens not considered to be human subjects research”.

### Human Platelet-rich plasma (PRP) collection

Platelet-rich plasma (PRP) samples were obtained from healthy donors through the Hoxworth Blood Center (HBC). PRP is routinely collected at HBC as part of standard platelet transfusion products, with each unit containing approximately 300 mL. The PRP used in this study was sourced from extra PRP allocated for quality control testing, performed independently by the HBC quality control team, who are not affiliated with this research. For each experiment, 2-5 mL of PRP was requested from the quality control team. Prior to transfer to our laboratory, all PRP samples underwent de-identification procedures to ensure donor anonymity. The consistent quality of these samples, maintained by the HBC quality control process, provided a reliable and reproducible source of human platelets for evaluating the albumin-platelet interaction.

### Platelet sample preparation

For each platelet force test, 20 µL of PRP was transferred to an Eppendorf tube and mixed with 0.5 mL of Buffer 1 (113 mM NaCl, 4.3 mM K_2_HPO_4_, 4.2 mM Na_2_HPO_4_, 24.3 mM NaH_2_PO_4_, 5.5 mM dextrose, pH 6.3), supplemented with 0.02 U/mL apyrase (M0398S, NEB Inc) and 1 µM prostaglandin E₁ (538903, Merck). The suspension was centrifuged at 800 × g for 2 minutes, the supernatant was removed, and the platelet pellet was resuspended in 0.5 mL of serum-free DMEM (30-2002, ATCC). Platelet concentration was determined using a hemocytometer and adjusted to levels required by the tests.

For platelet aggregation test, 0.3 mL of PRP was transferred to an Eppendorf tube and mixed with 0.5 mL of Buffer 1, supplemented with 0.02 U/mL apyrase (M0398S, NEB Inc) and 1 µM prostaglandin E₁ (538903, Merck). The suspension was centrifuged at 800 × g for 2 minutes, the supernatant was removed, and the platelet pellet was resuspended in 0.6 mL of serum-free DMEM (30-2002, ATCC).

Dependent on the tests, the platelets were either activated or maintained at resting state. To induce platelet activation, the platelet suspension was supplemented with 10 µM adenosine diphosphate (ADP, J60672.03, ThermoFisher Scientific) To maintain platelets in a resting state for activation assays, the suspension was instead supplemented with 0.02 U/mL apyrase to degrade residual ADP in the medium, which may activate platelets prematurely.

### Albumin-casein surface co-adsorption

Albumin and casein were adsorbed onto separate and adjacent regions of a glass surface. 100 µg/mL Alexa488-labeled human serum albumin (009-540-051, Jackson ImmunoResearch) and 100 µg/mL casein (37582, ThermoFisher Scientific) were prepared in phosphate buffered saline (PBS). 200 µL casein solution was added to the glass surface of a glass-bottomed Petri-dish (D35-14-1.5-N, Cellvis) which was subsequently incubated for 30 minutes at room temperature in a humidity chamber (a sealed box containing water-soaked cotton). The casein droplet was washed with copious PBS, and 200 µL albumin solution was added to the Petri-dish and cover the entire glass surface. The Petri-dish was incubated for another 30 minutes and washed by PBS. At this stage, casein was adsorbed in a circular region on the glass, and albumin was adsorbed on the entire surface except the casein-adsorbed region. Platelet adhesion was tested on this glass surface.

### Albumin-casein surface micropatterning

Protein micropatterning was performed using polydimethylsiloxane (PDMS) stamps. A PDMS stamp (15 µm diameter, 15 µm spacing, 15 µm height; Research Microstamps) was incubated with 1 mg/mL casein solution for 5 minutes at room temperature. The stamp surface was then rinsed with deionized water and dried using compressed air for approximately 5 seconds. The casein-coated stamp was gently placed onto the glass surface of a glass-bottom Petri dish (D35-14-1.5-N, Cellvis) and left in contact for 10 seconds. The stamp was then carefully removed to avoid lateral displacement that could smear the printed protein pattern. The patterned surface was rinsed with PBS, followed by incubation with 100 µg/mL Alexa488-labeled human serum albumin in PBS for 5 minutes. Finally, the surface was washed three times with PBS.

### Platelet activation and adhesion assay

Activated platelets or resting platelets at concentration of 4.0 × 10^4^/µL were seeded onto glass coverslips pre-coated with 10 µg/ml fibronectin (F1141, Sigma-Aldrich), 100 µg/mL fibrinogen (F3879, Sigma-Aldrich), 100 µg/mL ultrapure bovine serum albumin (AM2616, Invitrogen), 100 µg/mL ultrapure human serum albumin (A8763, Sigma-Aldrich), 100 µg/mL recombinant human serum albumin (HSA-H5220, Acro Biosystems) or 100 µg/ml casein, and incubated at 37 °C in a cell incubator containing 5% CO₂ for 30 minutes. Platelets were fixed and stained with phalloidin for F-actin imaging.

### HeLa cell adhesion assay

HeLa cells were cultured under standard conditions and prepared for adhesion assays as follows. The cells were detached with a mild detachment buffer [100 mL 10× HBSS, 10 mL 1 M HEPES, 10 mL 7.5% sodium bicarbonate, 2.4 mL 500 mM EDTA, and 878 mL ultrapure water; adjusted to pH 7.4]. Detached cells were resuspended in fresh culture medium and seeded onto glass coverslips pre-coated with fibronectin, fibrinogen, albumin, or casein and incubated at 37 °C in a cell incubator containing 5% CO₂ for 1 hours, followed by fixation, staining and imaging.

### Platelet aggregation assay

To test if denatured albumin mediate platelet aggregation, PRP-derived platelets were resuspended in serum-free cell culture medium at a concentration of 4.0 × 10^5^/µL, comparable to that in whole blood. The platelet sample was aliquoted into centrifuge tubes, each containing 0.6 mL of platelet solution. These platelet solutions were further supplemented with 50 µg/mL native HSA (A8763, Sigma-Aldrich), 50 µg/mL denatured HSA (heated at 90°C for 5 min), or 50 µg/mL fibrinogen, respectively. No ADP was added to these samples. The platelet samples in tubes were incubated at 37 °C on a shaker operating at 150 rpm (round per minute) for 30 minutes. The sample shaking is to emulate fluidic stress experienced by platelets. Afterwards, these platelet samples were plated onto glass-bottom Petri-dishes, incubated at static condition for 5 min, and then fixed and stained for imaging. The resulted platelet thrombi were imaged by a confocal microscope.

To examine the recruitment of denatured albumin and fibrinogen to platelet thrombi, in another experiment setting, platelet solutions were supplemented with 50 µg/mL heat-denatured HSA-Alexa488 (009-540-051, Jackson ImmunoResearch) or 50 µg/mL fibrinogen-Alexa488 (F13191, ThermoFisher Scientific), respectively. No ADP was added to these samples. The platelet samples after 30-min shaking were plated onto glass-bottom Petri-dishes, incubated at 37 °C for 5 minutes, and then fixed and stained for imaging. The resulted platelet thrombi, denatured HSA and fibriongen were imaged by a confocal microscope.

Platelet aggregation assays were also tested with albumin-Alexa488 denatured by 0.1% ethanol or 10 µM H_2_O_2_. For these tests, albumin-Alexa488 was denatured at a concentration of 5 mg/mL, supplemented with 0.1% ethanol or 10 µM H_2_O_2_. With two-hour denaturation process, the denatured albumin (dnAlb) was added to platelet solutions without removing ethanol or H_2_O_2_, which were diluted by 100-fold, resulting in 50 µg/mL dnAlb, 0.001% ethanol or 0.1 µM H_2_O_2_ in the platelet solutions. The low levels of ethanol or H_2_O_2_ were not expected to significantly impact platelet functions. In the control test of platelet aggregation using native albumin-Alexa488, the albumin did not go through denaturation process, and 0.001% ethanol and 0.1 µM H_2_O_2_ were added to the platelet solution.

### DNA strands for ITS synthesis

DNA oligonucleotides were ordered from Integrated DNA Technologies, Inc. The following modifications were used: /5DBCO/ denotes a DBCO group at the 5’ end, which is used to react with an azide group through copper-free click chemistry; /3BHQ_2/ refers to the Black Hole Quencher 2 at the 3’ end; /5ThioMC6-D/ denotes a 5’ thiol modification with a six-carbon spacer, which after reduction, reacts with a maleimide group; /5BiosG/ denotes a biotin tag at the 5’ end; /iAtto647NN/ and /5Atto647NN/ indicate conjugation of the Atto647N fluorescent dye at internal or 5’ positions, respectively; /iAmMC6T/ represents an internal thymine (dT) modified with a C6 amine spacer. This amine group was used for biotinylation by reacting with NHS-PEG12-Biotin (A35389, Thermo Fisher Scientific). /3BioTEG/ represents a biotin tag conjugated to the 3’ end of the DNA through a trimer of ethylene glycol.

### DNA strands after conjugating with ligand or biotin

To synthesize Alb-ITS and RGD-ITS with desired tension tolerance (*T*_tol_) values, DNA strands 1, 2, 4-6 were conjugated with Albumin, RGD peptide or biotin according to protocols included in the content below. After the conjugation, DNA strands are shown below:

### Preparing azide-labeled human serum albumin (HSA-azide)

6.7 mg of human serum albumin (A8753, Sigma-Aldrich, purity ≥99%) was dissolved in 1 mL of PBS (PH 7.2) to obtain a 100 µM albumin solution. A 75 mM Azide-PEG3-maleimide solution (CCT-AZ107, Vector Laboratories) in DMSO was prepared following the manufacturer’s instruction. 14 µL of the 75 mM Azide-PEG3-maleimide solution was added to the 1 mL albumin solution, yielding a final concentration of 1 mM Azide-PEG3-maleimide, corresponding to a 1:10 molar ratio of albumin to Azide-PEG3-maleimide. The maleimide group reacts with the thiol of a single cysteine residue in human serum albumin (HSA). After reacting for 1 hour at room temperature, the albumin-azide solution was purified by passing it through desalting columns (89492, Thermo Scientific) four times to remove unreacted azide-PEG3-maleimide. The desalting columns were thoroughly equilibrated with PBS by buffer exchange before use. The buffer exchange is expected to remove the sodium azide in the columns. At this stage, the albumin was conjugated with azide, producing HSA-azide which is ready for following DNA conjugation or biotin conjugation.

HSA can be functionalized with one single azide, owing to the fact that HSA has a single reactive thiol group in its molecular structure. Although HSA contains 35 cysteines, 34 form 17 disulfide bonds, leaving only one free cysteine (Cys34) with a reactive thiol group. Cys34 is also located at the outer domain of the albumin protein, making it sterically accessible. This allows the conjugation of a single azide moiety, and subsequently a single DNA molecule or a single biotin tag, to each HSA molecule. Refer to Supplementary Figs 6 and 7.

The pH of PBS was adjusted to 7.2 to promote the maleimide-thiol reaction while minimizing the maleimide-amine reaction, which could undesirably label albumin with additional azide groups via lysine residues.

### Conjugating albumin with the DNA upper strand

To conjugate a single-stranded DNA to the albumin, 52.5 µL × 1 mM DNA 1 in PBS (Table 1) was added to the 1 mL × 100 µM HSA-azide solution. The amounts of DNA and HSA-azide can be scaled down while maintaining the same molar ratio. The solution was incubated for 8 h at room temperature and overnight at 4 °C. The albumin and DNA were conjugated through DBCO-azide reaction, a copper-free click chemical reaction, producing the DNA 1-Alb in Table 2 at a concentration of 50 µM. No further purification is needed as the excessive unreacted albumin will be eventually washed during Alb-ITS surface attachment. This purification-free protocol minimizes the unintended albumin denaturation during albumin-DNA conjugation. The albumin-DNA conjugate is hybridized with complementary DNA to synthesize Alb-ITS. Extra albumin-DNA conjugate was stored at-20°C.

**Table 1.**
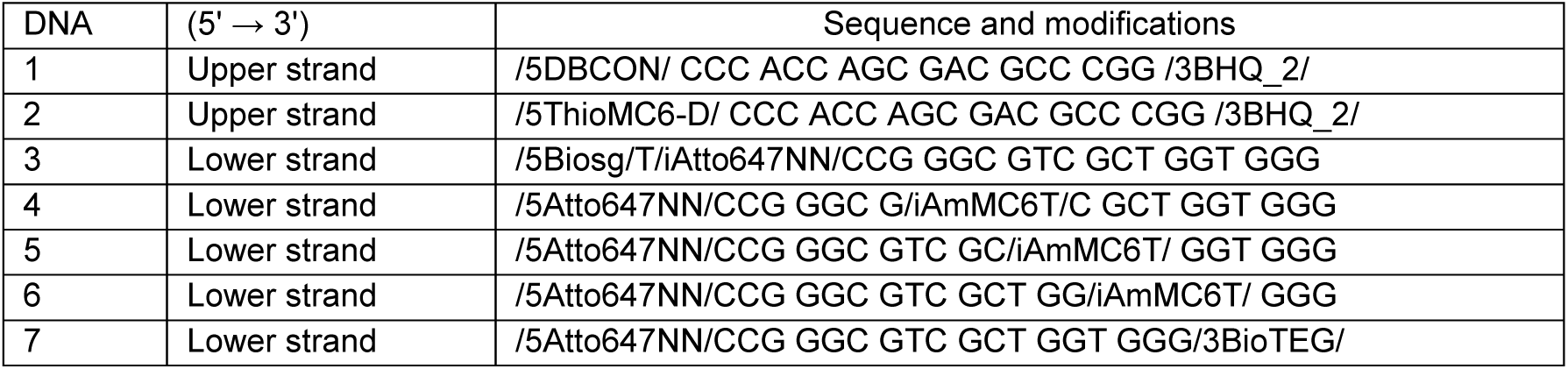
DNA constructs ordered for ITS synthesis.

**Table 2.**
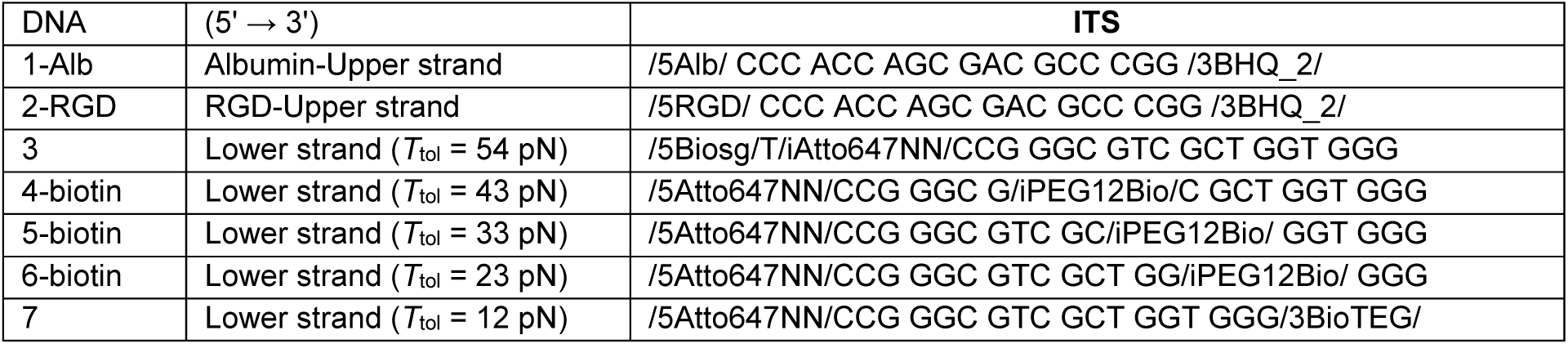
DNA conjugated with ligands or biotin. Ready for ITS synthesis.

The albumin-DNA conjugation can be verified by SDS-PAGE, as shown in Supplementary Fig. 7. Typically, further purification is unnecessary. However, if a substantial fraction of HSA is labeled with more than one DNA molecule, as indicated by multiple bands on the protein gel, additional purification can be performed using FPLC (Fast Protein Liquid Chromatography) to isolate HSA conjugated with a single DNA strand.

### Conjugating albumin with a single biotin

Monobiotinylated albumin was prepared by reacting HSA-azide with Biotin-PEG4-DBCO (CCT-A105-5, Vector Laboratories). HSA-azide was prepared at a concentration of 100 µM in PBS. Biotin-PEG4-DBCO was prepared at 10 mM in DMSO. These two reactants were mixed at a molar ratio of 1:10 and reacted for 2 h at room temperature. The product was dialyzed with the kit (PURN12010, Sigma-Aldrich), resulting HSA-biotin (Alb-biotin) which was used in subsequent experiments.

### Conjugating RGD to the DNA upper strand

Integrin peptide ligand RGD (Arginine-Glycine-Aspartate) was conjugated to the upper DNA strand of ITS constructs using this protocol. A reducing solution containing 500 mM TCEP (77720, ThermoFisher Scientific) and 500 mM EDTA (AM9260G, Invitrogen) in PBS (phosphate-buffered saline) was prepared. This solution was used to reduce the disulfide bond in the /5ThioMC6-D/ modification on the DNA strand, converting it into a free thiol group for subsequent thiol-maleimide conjugation. To perform the reduction, 10 μL of the TCEP/EDTA solution was added to 100 μL of 2 mM DNA 2 in PBS (Table 1). The mixture was incubated at room temperature for 30 minutes (referred to as Solution 1).

In parallel, 10 mg of RGD-NH₂ (PCI-3696-PI, Biosynth) was dissolved in 200 μL of ultrapure water. This solution was then added directly to 2 mg of solid sulfo-SMCC (A39268, Thermo Fisher Scientific) in a vial dipped in an ultrasonic bath to facilitate rapid dissolution. Subsequently, 22 μL of 10× PBS was added to adjust the mixture to a final concentration of 1× PBS. The solution was incubated at room temperature for 10 minutes (referred to as Solution 2). Solution 1 and 2 were then mixed and incubated for 1 h at room temperature and overnight at 4 °C. RGD is now conjugated to the DNA through sulfo-SMCC which is a heterolinker with a maleimide (reacting with the thiol of the DNA) at one end and an NHS ester (reacting with the amine of RGD-NH_2_) at the other end. The unreacted SMCC, RGD and TCEP were removed via ethanol precipitation. The precipitated RGD-DNA was re-dissolved in PBS, and the concentration was adjusted to 100 μM using a Nanodrop 2000 Spectrophotometer (ND-2000, Thermo Fisher Scientific), producing the DNA 2-RGD in the Table 2.

### Conjugating PEG12-biotin to the lower DNA strands (DNAs 4-6 in the Table 1)

For ITS constructs with intermediate *T*_tol_ values, the biotin was conjugated to a thymine modified with an amine in the middle of a lower-strand indicated by/iAmMC6T/. During conjugation, 100 μM×200 μL DNA in PBS was added to one vial containing 1 mg NHS-PEG12-Biotin (A35389, no-Weight format, Thermo Fisher Scientific). Let the mixture react for 2 h at the room temperature. Ethanol precipitation was conducted to purify the DNA conjugated with PEG12-Biotin. The biotin-labeled DNA was re-dissolved in PBS and its concentration was adjusted to 100 μM (DNAs 4-biotin, 5-biotin and 6-biotin in Table 2).

### Alb-ITS and RGD-ITS assembly and storage

Alb-ITS constructs were assembled by hybridizing the upper DNA strand (DNA 1-Alb in Table 2) with complementary lower DNA strands (DNAs 3-7 in Table 2). The resulting *T*_tol_ values are shown in Table 2. The strands were mixed at a molar ratio of 1.2:1 (upper:lower) to ensure complete hybridization of the dye-labeled lower strands with the quencher-labeled upper strand, and the final concentration of double-stranded DNA was adjusted to 10 µM. The mixtures were incubated at room temperature for 2 h and at 4 °C overnight. DNA annealing was omitted to preserve the structural integrity of albumin. Hybridization of the 18 base-paired dsDNA generally does not require annealing either.

After synthesis, Alb-ITS should be aliquoted and stored at-20 °C. Samples stored at 4 °C were observed to gradually denature within 1-2 weeks. Each frozen aliquot should be used only once to avoid multiple freeze-thaw cycles.

RGD-ITS constructs were prepared similarly, using DNA 2 as the upper strand and DNAs 3-7 as the lower strands. DNA annealing was also omitted to maintain consistency with the Alb-ITS preparation. RGD-ITS can be stored at either 4°C or-20°C without apparent quality change after months.

### Denaturing albumin, albumin-biotin, albumin-Alexa488 or Alb-ITS by heating

To induce albumin denaturation by heat, albumin products (1 mg/mL) or Alb-ITS (10 µM) in 1.5 mL centrifuging tubes with at least 20 µL volume was placed in a heating block at designed temperature for designed durations. The samples were then cooled to room temperature and were ready for further test.

### Denaturing albumin or Alb-ITS using SDS, ethanol or hydrogen peroxide

Alb-ITS were also denatured using sodium dodecyl sulfate (SDS), ethanol, or hydrogen peroxide. Alb-ITS (10 µM) solutions were treated with these reagents at the concentrations and durations specified in the manuscript. During platelet force assays, denatured Alb-ITS (dnAlb-ITS) were diluted 100-fold to a final concentration of 0.1 µM, along with corresponding dilutions of the denaturing reagents. These diluted chemicals did not noticeably affect the biotin-avidin interaction responsible for tethering dnAlb-ITS to the surface.

### Preparing biotin-functionalized glass surfaces

ITS constructs were immobilized on a surface via biotin-avidin interactions, requiring the surface to be biotin-functionalized for the immobilization of avidin. On top of the immobilized avidin, biotin-tagged ITS constructs were tethered.

For biotin-functionalization, glass coverslips placed in a jar were cleaned by sequential sonication in Milli-Q water for 15 min, acetone for 15 min, and rinsed three times with Milli-Q water. The cleaned coverslips were then sonicated in 1 M KOH (potassium hydroxide) for 45 min, followed by three rinses with water and three rinses with methanol. Amino-silanization was performed on these coverslips to introduce amine groups on the surface, which were subsequently used to immobilize biotin groups via an amine-NHS ester reaction.

For amino-silanization, the jar containing coverslips was filled with a solution prepared by mixing 150 mL methanol, 7.5 mL acetic acid, and 1.5 mL (3-aminopropyl) triethoxysilane (APTES; 741442, Sigma-Aldrich). During silanization, APTES grafted amine groups onto the glass surface. To introduce biotin groups, a 70 µL droplet of 5 mg/mL sulfo-NHS-biotin (21217, Thermo Scientific) in PBS was placed on the amine-functionalized coverslip and sandwiched with a second coverslip. Biotinylation occurred via the amine-NHS ester coupling reaction. After incubation at the room temperature for 2 h, these coverslips were separated, washed with copious pure water and dried with compressed air.

For platelet force assays, the biotin-functionalized coverslips were cut to appropriate sizes and affixed to bottomless Petri dishes (D35-14, Cellvis) using UV-curable resin. Excess coverslips were stored at-20 °C.

### Preparing PEGylated glass surfaces

Following the procedure listed in “Preparing biotin-functionalized glass surfaces”, after the step of silanization which grafts amine groups onto the glass surface, a 70 µL droplet of PEG solution, instead of sulfo-NHS-biotin solution, was placed on the amine-functionalized coverslip and sandwiched with a second coverslip. The remaining procedure is the same as preparing for biotin-functionalized glass surfaces.

The PEG mixture consisted of 20 mg Biotin-PEG (Biotin-PEG-SVA, 5 kDa, Laysan Bio) and 200 mg mPEG (mPEG-SVA, 5 kDa, Laysan Bio). At this concentration, the PEG solution is viscous. To ensure proper dispersion and dissolution, vortex the solution and then centrifuge it for 2 min at 10,000 × g. The PEG solution should be used immediately after preparation, as the NHS ester groups in the PEG polymers hydrolyze rapidly and may lose reactivity within minutes.

### Tethering Alb-ITS/dnAlb-ITS/RGD-ITS on biotin-functionalized surfaces

On the Petri-dish assembled with a biotin-functionalized coverslip, 100 µg/ml neutravidin (31000, Thermo Fisher Scientific) was added and incubated for 30 minutes at 4 °C to coat neutravidin on the biotin-presenting surface. The coverslip was rinsed by PBS three times, and then incubated with 0.1 µM Alb-ITS, dnAlb-ITS or RGD-ITS for 30 min at 4 °C. Each ITS construct has a biotin tag, enabling the surface-tethering of the ITS on the glass surface. The surface was rinsed by PBS three times and ready for cell plating. This sequential assembly results in a stable immobilization configuration: glass-biotin → neutravidin → biotin-tagged ITS. Neutravidin, with its four biotin-binding sites, ensures strong tethering of the ITS on the surface.

### Platelet force assays on ITS surfaces

Platelets at concentration of 4.0 × 10^4^/µL in serum-free medium were activated by 10 µM ADP and seeded onto the ITS-tethered surfaces. The samples were incubated at 37 °C in a cell incubator containing 5% CO₂ for 30 minutes. Platelets were fixed and stained with phalloidin or antibodies. The platelet force signals and cell structural signals were imaged by a total internal reflection fluorescence (TIRF) microscope.

### Platelet force Assays with inhibitors of integrin or myosin

For the inhibition of integrin α_IIb_β_3_, before cell plating, ADP-activated platelet samples were supplemented with 20 µg/mL Abciximab (MA5-47865, ThermoFisher Scientific catalog #, company), 5 µg/mL eptifibatide (SML1042-10MG, Sigma-Aldrich catalog #, company), or 5 µg/mL Tirofiban (50-257-7097, Fisher Scientific). For myosin II inhibition, 20 µM blebbistatin (B0560, ThermoFisher Scientific) was added to the platelet samples. The treated platelets samples were plated on either Alb-ITS or RGD-ITS surfaces as indicated in the manuscript. Platelets on these surfaces were incubated at 37 °C in an incubator with 5% CO₂ for 30 minutes, followed by fixation, staining and imaging.

### Immunostaining

For the co-imaging of protein structure and dnAlb-transmitted forces in platelets, immunostaining was performed over platelets incubated and fixed on dnAlb-ITS surfaces. Platelets were seeded onto dnAlb-ITS surfaces and incubated for 30 minutes at 37 °C. Following incubation, cells were fixed with 4% paraformaldehyde and permeabilized with 0.1% Triton X-100. After blocking with 1% casein for 1 hour, platelet samples were washed three times with PBS, allowing a 5-minute interval between each wash. To visualize structural protein actin filaments, cells were stained with phalloidin conjugated with Alexa488 (A12379, Invitrogen). Samples were incubated with 0.1 µM phalloidin for 30 minutes at room temperature, then washed three times with PBS. To stain target proteins, cell samples were treated with 5 µg/mL primary antibodies against integrin β_1_ (14-0299-82, Thermo Fisher Scientific), integrin β_3_ (14-0619-82, Thermo Fisher Scientific), vinculin (FAK100, Sigma-Aldrich) or p-MLC 2 (3675T, Cell signaling technology) for 1 hour at room temperature. After incubation and three-time wash, cells were incubated with the secondary antibody, Alexa-488 goat anti-mouse IgG (A-32723, Invitrogen), for 1.5 hours at room temperature. Finally, cells were washed three times with PBS washes to remove unbound stain protein and visualized with Alb-ITS-mediated force transmission using TIRF microscopy.

### Microscopy and data processing

Imaging was conducted using either a Nikon Ti-2E Confocal microscope or a Nikon Ti-2E TIRF microscope (total internal reflection fluorescence), depending on the specific requirements of the experiment. For real-time imaging, a lens heater was used to support cell viability. All imaging was acquired with the 100× oil immersion objective lens. Laser beams at 405, 488, 561 and 640 nm wavelengths were used as the excitation light sources. Image processing and analysis were conducted using the software (NIS-element) provided with the microscope combined with customized Matlab codes. MatLab and Graphpad prism software were used for graph plotting.

### Quantification and Statistical Analysis

In the manuscript, measurements are reported as mean ± standard deviation of the mean (SD). The results of key experiments were repeated three times and data were pooled together for statistical analysis. The volume of sample analyzed is specified in the figure legends. P values in figures were obtained using two-tailed unpaired t-test with Welch’s correction using Graphpad prism.

## DATA AVAILABILITY

All data supporting the findings of this study are included within the article and its supplementary files. For any further information, inquiries can be directed to the corresponding author, who will respond to and fulfill such requests.

## ACKNOWLEDGMENTS

This research was supported by the research fund from National Institute of General Medical Sciences (R35GM128747, X.W.) and National Science Foundation (grant 2204447 to L.Q. and X.W.). We are grateful to the quality control team (Stacy Braun, Katheryn Boedecker, Ciara Combs, Sheryl Heeb, Lindsey Marquez, Miriam Michael, Kimberly Molumby, Priddy Taylor, Lindsey O’Bannion, Amy Parker, and Jennifer O’Connor) at Hoxworth blood center for providing platelet rich plasma with consistent sample quality.

## AUTHOR CONTRIBUTIONS

X. W. conceived the research project; X.W. and V.P. designed experiments; V. P. performed experiments, with contributions from S. K., S. J. W., A. P., and L. Q.; V.P. and X. W. analyzed results and plotted the graphs. X. W. and V. P. wrote the manuscript.

## REFERENCES

1. Farrugia, A. Albumin usage in clinical medicine: tradition or therapeutic? Transfus Med Rev 24, 53–63 (2010).

2. Bellier, J.P. et al. Identification of fibrinogen as a plasma protein binding partner for lecanemab biosimilar IgG. Ann Clin Transl Neur 11, 3192–3204 (2024).

3. Furie, B. & Furie, B.C. The Molecular-Basis of Blood-Coagulation. Cell 53, 505–518 (1988).

4. Coller, B.S. & Shattil, S.J. The GPIIb/IIIa (integrin αIIbβ3) odyssey: a technology-driven saga of a receptor with twists, turns, and even a bend. *Blood*, The Journal of the American Society of Hematology 112, 3011–3025 (2008).

5. Nieswandt, B. et al. Glycoprotein VI but not α2β1 integrin is essential for platelet interaction with collagen. The EMBO journal (2001).

6. Nieswandt, B. & Watson, S.P. Platelet-collagen interaction: is GPVI the central receptor? Blood 102, 449–461 (2003).

7. Levy, J.H., Szlam, F., Tanaka, K.A. & Sniecienski, R.M. Fibrinogen and hemostasis: a primary hemostatic target for the management of acquired bleeding. Anesth Analg 114, 261–274 (2012).

8. Kattula, S., Byrnes, J.R. & Wolberg, A.S. Fibrinogen and Fibrin in Hemostasis and Thrombosis. Arterioscler Thromb Vasc Biol 37, e13–e21 (2017).

9. Singh-Zocchi, M., Andreasen, A. & Zocchi, G. Osmotic pressure contribution of albumin to colloidal interactions. P Natl Acad Sci USA 96, 6711–6715 (1999).

10. Merlot, A.M., Kalinowski, D.S. & Richardson, D.R. Unraveling the mysteries of serum albumin-more than just a serum protein. Front Physiol 5 (2014).

11. Amiji, M., Park, H. & Park, K. Study on the prevention of surface-induced platelet activation by albumin coating. J Biomater Sci Polym Ed 3, 375–388 (1992).

12. Gaertner, F. et al. Migrating Platelets Are Mechano-scavengers that Collect and Bundle Bacteria. Cell 171, 1368–1382 e1323 (2017).

13. Park, K., Mao, F.W. & Park, H. The minimum surface fibrinogen concentration necessary for platelet activation on dimethyldichlorosilane-coated glass. J Biomed Mater Res 25, 407–420 (1991).

14. McCarty, O. et al. Evaluation of the role of platelet integrins in fibronectin-dependent spreading and adhesion. Journal of thrombosis and haemostasis 2, 1823–1833 (2004).

15. Goodman, S.L., Cooper, S.L. & Albrecht, R.M. The effects of substrate-adsorbed albumin on platelet spreading. J Biomater Sci Polym Ed 2, 147–159 (1991).

16. Sivaraman, B. & Latour, R.A. The adherence of platelets to adsorbed albumin by receptor-mediated recognition of binding sites exposed by adsorption-induced unfolding. Biomaterials 31, 1036–1044 (2010).

17. Hylton, D.M., Shalaby, S.W. & Latour, R.J.r. Direct correlation between adsorption-induced changes in protein structure and platelet adhesion. J Biomed Mater Res A 73a, 349–358 (2005).

18. Handagama, P., Scarborough, R.M., Shuman, M.A. & Bainton, D.F. Endocytosis of Fibrinogen into Megakaryocyte and Platelet Alpha-Granules Is Mediated by Alpha(Iib)Beta(3) (Glycoprotein-Iib-Iiia). Blood 82, 135–138 (1993).

19. Sakurai, Y. et al. Platelet geometry sensing spatially regulates alpha-granule secretion to enable matrix self-deposition. Blood 126, 531–538 (2015).

20. Vogt, R.F., Phillips, D.L., Henderson, L.O., Whitfield, W. & Spierto, F.W. Quantitative Differences among Various Proteins as Blocking-Agents for Elisa Microtiter Plates. J Immunol Methods 101, 43–50 (1987).

21. Cho, J. & Mosher, D.F. Role of fibronectin assembly in platelet thrombus formation. Journal of Thrombosis and Haemostasis 4, 1461–1469 (2006).

22. Zaidi, T.N., McIntire, L.V., Farrell, D.H. & Thiagarajan, P. Adhesion of platelets to surface-bound fibrinogen under flow. Blood 88, 2967–2972 (1996).

23. Heggestad, J.T., Fontes, C.M., Joh, D.Y., Hucknall, A.M. & Chilkoti, A. In Pursuit of Zero 2.0: Recent Developments in Nonfouling Polymer Brushes for Immunoassays. Adv Mater 32 (2020).

24. Kapp, T.G. et al. A Comprehensive Evaluation of the Activity and Selectivity Profile of Ligands for RGD-binding Integrins. Sci Rep-Uk 7 (2017).

25. Rana, A., Westein, E., Niego, B. & Hagemeyer, C.E. Shear-Dependent Platelet Aggregation: Mechanisms and Therapeutic Opportunities. Front Cardiovasc Med 6, 141 (2019).

26. Kundu, S., Pal, K., Pyne, A. & Wang, X.F. Force-bearing phagocytic adhesion rings mediate the phagocytosis of surface-bound particles. Nat Commun 16 (2025).

27. Tu, Y., Pal, K., Austin, J. & Wang, X.F. Filopodial adhesive force in discrete nodes revealed by integrin molecular tension imaging. Curr Biol 32, 4386–4396 (2022).

28. Wang, Y.L. et al. Force-activatable biosensor enables single platelet force mapping directly by fluorescence imaging. Biosens Bioelectron 100, 192–200 (2018).

29. Kundu, S., Pandey, V., Pyne, A., Que, L. & Wang, X. Characterizing Integrin Tensions during Platelet Adhesion and Stiffness Sensing. Nano Lett 25, 11156–11163 (2025).

30. Wang, X.F. & Ha, T. Defining Single Molecular Forces Required to Activate Integrin and Notch Signaling. Science 340, 991–994 (2013).

31. Hashemzadeh, M., Furukawa, M., Goldsberry, S. & Movahed, M.R. Chemical structures and mode of action of intravenous glycoprotein IIb/IIIa receptor blockers: A review. Exp Clin Cardiol 13, 192–197 (2008).

32. Deibele, A.J. et al. Intracoronary eptifibatide bolus administration during percutaneous coronary revascularization for acute coronary syndromes with evaluation of platelet glycoprotein IIb/IIIa receptor occupancy and platelet function: the Intracoronary Eptifibatide (ICE) Trial. Circulation 121, 784–791 (2010).

33. Tam, S.H., Sassoli, P.M., Jordan, R.E. & Nakada, M.T. Abciximab (ReoPro, chimeric 7E3 Fab) demonstrates equivalent affinity and functional blockade of glycoprotein IIb/IIIa and alpha(v)beta3 integrins. Circulation 98, 1085–1091 (1998).

34. Bennett, J.S. Structure and function of the platelet integrin α IIb β 3. The Journal of clinical investigation 115, 3363–3369 (2005).

35. Bledzka, K., Qin, J. & Plow, E.F. Integrin αIIbβ3. Platelets, 227–241 (2019).

36. Lam, W.A. et al. Mechanics and contraction dynamics of single platelets and implications for clot stiffening. Nat Mater 10, 61–66 (2011).

37. Austin, J., Tu, Y., Pal, K. & Wang, X.F. Vinculin transmits high-level integrin tensions that are dispensable for focal adhesion formation. Biophys J 122, 156–167 (2023).

38. Lapham, S.C. The limits of tolerance: convicted alcohol-impaired drivers share experiences driving under the influence. Perm J 14, 26–30 (2010).

39. Reichmann, D., Voth, W. & Jakob, U. Maintaining a Healthy Proteome during Oxidative Stress. Mol Cell 69, 203–213 (2018).

40. Burgoyne, J.R., Oka, S., Ale-Agha, N. & Eaton, P. Hydrogen Peroxide Sensing and Signaling by Protein Kinases in the Cardiovascular System. Antioxid Redox Sign 18, 1042–1052 (2013).

41. Forman, H.J., Bernardo, A. & Davies, K.J.A. What is the concentration of hydrogen peroxide in blood and plasma? Arch Biochem Biophys 603, 48–53 (2016).

42. Erana, H. et al. A Protein Misfolding Shaking Amplification-based method for the spontaneous generation of hundreds of bona fide prions. Nat Commun 15, 2112 (2024).

43. Graf, P.C. et al. Activation of the redox-regulated chaperone Hsp33 by domain unfolding. J Biol Chem 279, 20529–20538 (2004).

44. Mishra, V. & Heath, R.J. Structural and Biochemical Features of Human Serum Albumin Essential for Eukaryotic Cell Culture. Int J Mol Sci 22 (2021).

45. Bito, R. et al. Degradation of oxidative stress-induced denatured albumin in rat liver endothelial cells. Am J Physiol-Cell Ph 289, C531–C542 (2005).

46. Paar, M., et al. Albumin in patients with liver disease shows an altered conformation. Commun Biol 4 (2021).

47. Tsao, F.H.C. & Meyer, K.C. Human Serum Albumin Misfolding in Aging and Disease. Int J Mol Sci 23 (2022).

48. Walter, E.J., Hanna-Jumma, S., Carraretto, M. & Forni, L. The pathophysiological basis and consequences of fever. Crit Care 20, 200 (2016).

49. Baler, K. et al. Electrostatic unfolding and interactions of albumin driven by pH changes: a molecular dynamics study. J Phys Chem B 118, 921–930 (2014).

50. Sharma, S.K., Goloubinoff, P. & Christen, P. Heavy metal ions are potent inhibitors of protein folding. Biochem Biophys Res Commun 372, 341–345 (2008).

