## Supplementary Figures for "Denatured Albumin Gains a Function of Regulating Platelet Activity"

### Supplemental Figures

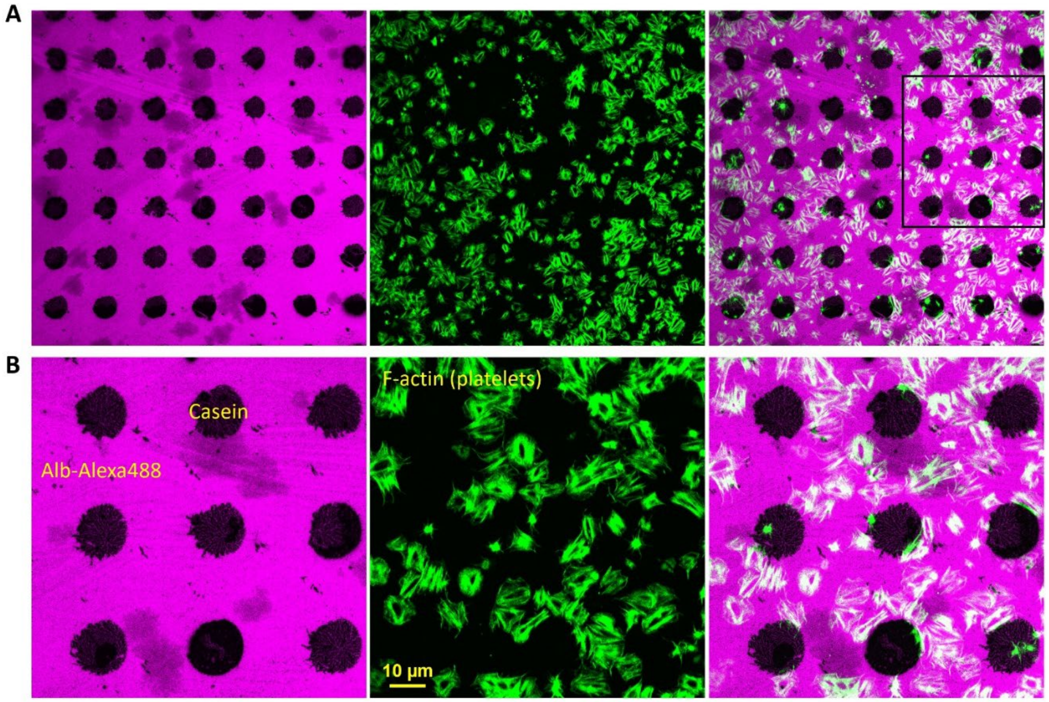


**Fig. S1. Platelets adhered on albumin-coated regions but not on casein-coated regions.**

(**A**) Platelets only adhered on albumin-coated regions. Casein was micropatterned onto the substrate using a PDMS stamp to generate a cluster of micro-islets (dark regions). The surface was subsequently incubated with Alexa488-labeled albumin (magenta color). Casein coatings effectively excluded albumin-Alexa488 from the islet regions, thereby restricting albumin adsorption to the background areas. (**B**) Zoom-in regions of (A). Platelets (green) were shown to adhere and spread only on albumin-adsorbed region (magenta). Platelets were fixed and stained with phalloidin for the cell imaging.

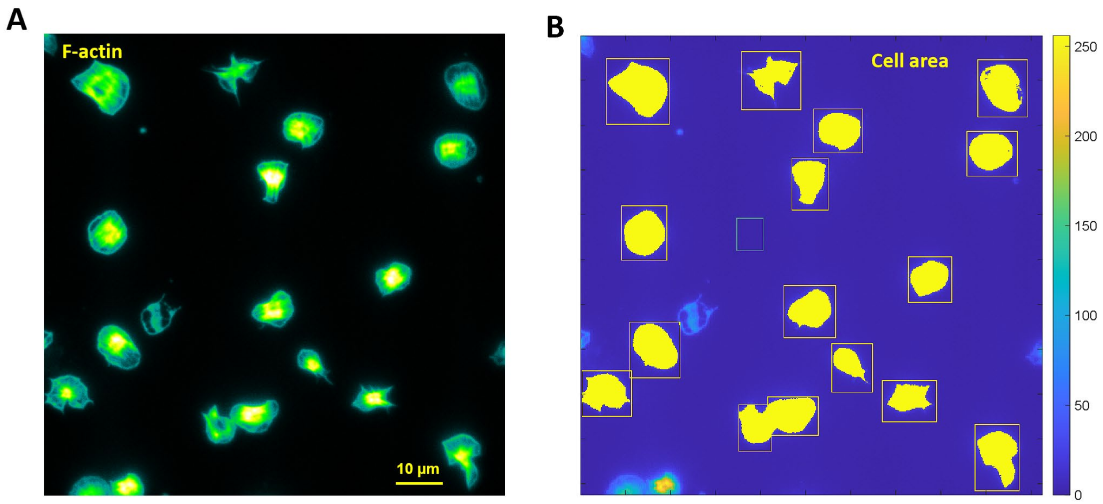


**Fig. S2. Algorithm for quantification of cell adhesion area.**

A MATLAB code was developed to analyze cell spreading areas. Each cell was manually selected using a rectangular region. The cell area was computed by counting the number of pixels with grayscale values exceeding a predefined threshold in the F-actin image. Each pixel is corresponding to an area: $0.16^2$ $\mu m^2$. (**A**) F-actin images of adherent platelets. (**B**) A sample of data processing which produces a list of platelet spreading areas. The regions highlighted in yellow represent platelet spreading areas.

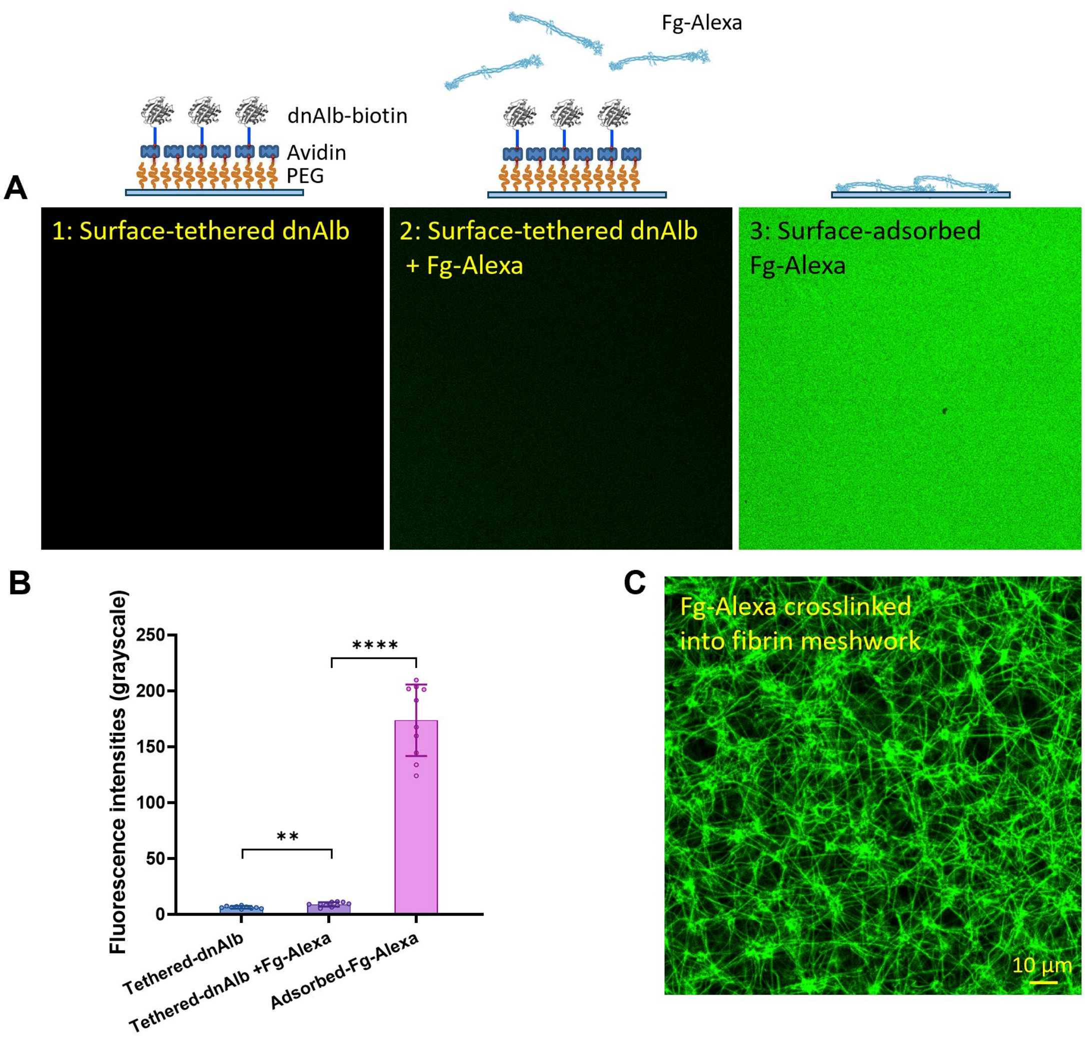


**Fig. S3. dnAlb has minimal or no binding capacity with fibrinogen.**

(**A**) Fluorescence images in the channel of Alexa488 over three surfaces prepared to verify the potential dnAlb-fibrinogen binding. Surface 1: dnAlb-biotin (100 µg/mL) was tethered on a PEGylated glass surface through the biotin-neutravidin interaction. Surface 2: On the dnAlb-tethered PEG surface, fibrinogen-Alexa488 (Fg-Alexa, 20 µg/mL) was added and incubated for 30 min. The surface was rinsed by PBS three times. Surface 3: Fg-Alexa (20 µg/mL) was coated on a glass surface through physical adsorption. After 30 min incubation, the surface was rinsed by PBS three times. (**B**) Fluorescence intensities on the three surfaces. Each data point represents the grayscale value per pixel averaged over one fluorescent image; N = 10 images for each surface. Fluorescence intensities on these surfaces are 6.4±1.1 (surface 1), 8.9±2.0 (surface 2) and 173.7±32.0 (surface 3), respectively. The slight increase in fluorescence from surface 1 to surface 2 indicates that dnAlb has minimal or no binding capacity with fibrinogen. This slight fluorescence increase may result from fibrinogen deposition on the PEGylated glass, which does not completely prevent protein adsorption. (**C**) The identity and function of Fg-Alexa were validated by examining its ability to form fibrin meshwork. Fg-Alexa was mixed with platelet-rich plasma (PRP), and thrombin (2 U/mL) was added to catalyze fibrinogen polymerization. A well-defined meshwork was observed in the Alexa488 fluorescence channel, confirming that Fg-Alexa retained fibrinogen functionality.

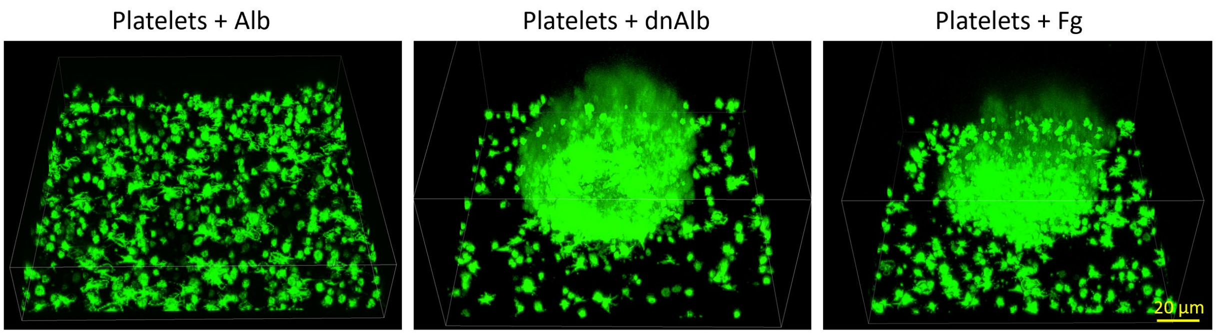


**Fig. S4. Three-dimensional views of thrombi formed in platelet solutions supplemented with Alb, dnAlb or fibrinogen.**

Platelets were purified from PRP and re-suspended in serum-free medium at a concentration of $4\times10^5$/µL. No ADP was added. The suspensions were supplemented with 50 µg/mL human serum albumin (Alb), 50 µg/mL denatured human serum albumin (dnAlb), or 50 µg/mL fibrinogen (Fg), respectively. These platelet solutions were incubated at 37 °C for 30 min on a shaker operating at 150 rpm (emulating fluidic stress), and then transferred to glass-bottom Petri dishes. After an additional 5 min of incubation, the platelets were fixed and stained for imaging. Platelet aggregates (thrombi) were commonly observed in dnAlb- and Fg-supplemented samples but were absent in those supplemented with Alb.

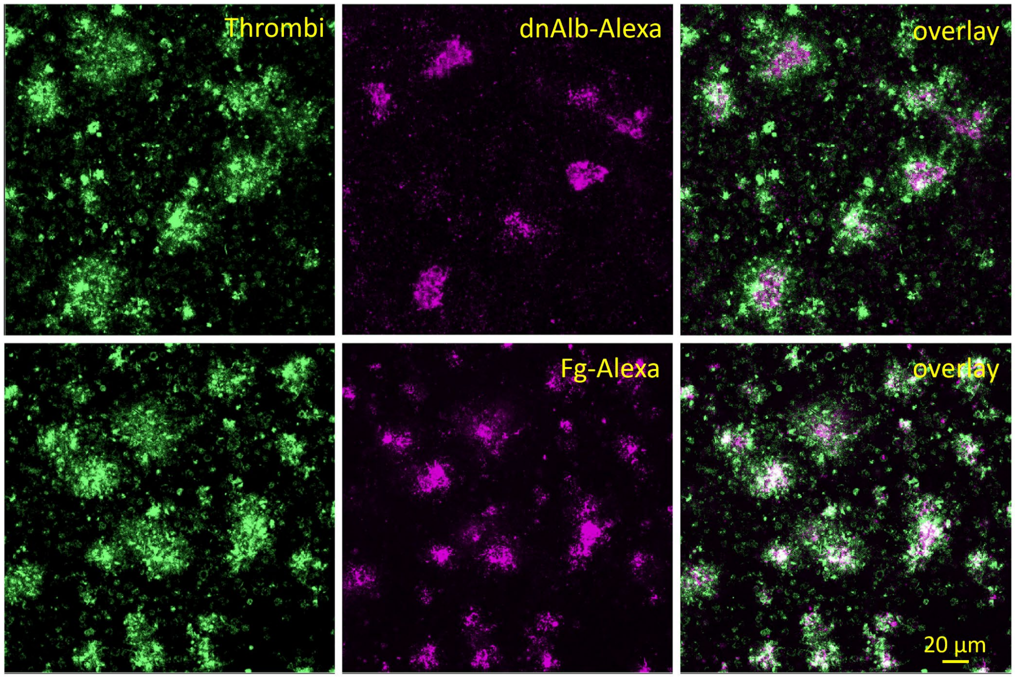


**Fig. S5. Recruitment of denatured albumin(dnAlb) or fibrinogen (Fg) to thrombi.**

Platelets were resuspended in serum-free cell culture medium at a concentration of $4.0 \times 10^5/\mu L$. The platelet solutions were further supplemented with 50 µg/mL denatured albumin-Alexa488 (dnAlb-Alexa) or 50 µg/mL fibrinogen-Alexa488 (Fg-Alexa), respectively. These platelet solutions were loaded onto glass-bottom Petri dishes, and incubated at 37 °C with shaking (150 rpm) on a shaker for 30 minutes. The recruitment of the dye-labeled proteins to the platelet aggregates (thrombi) was imaged with confocal microscopy.

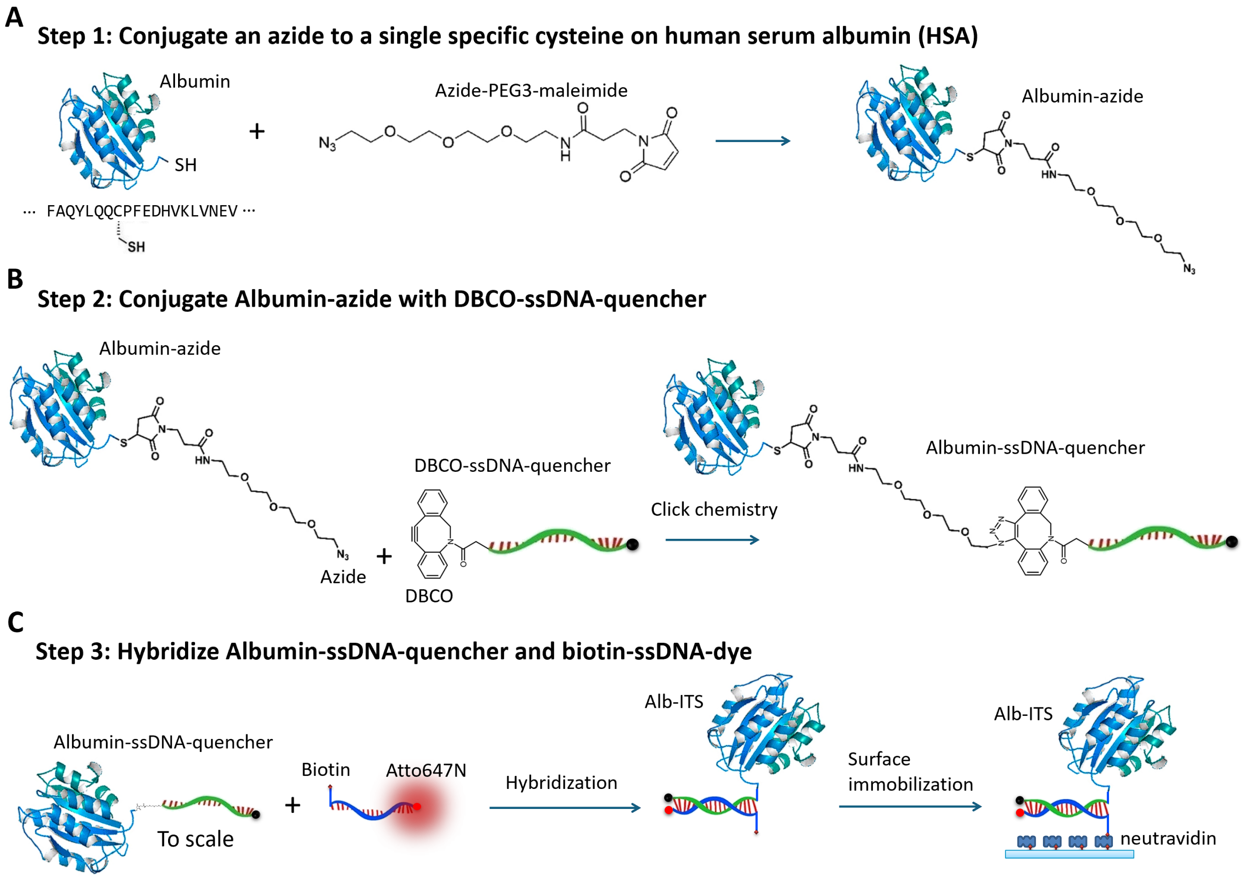


**Fig. S6. Synthesis of albumin-ITS.**

Three steps to synthesize the tension sensor equipped with albumin. (**A**) A single azide is conjugated to a specific cysteine residual in human serum albumin. Refer to Materials and Methods. (**B**) The HSA-azide was conjugated with DBCO-ssDNA-quencher through copper-free click chemistry. The reaction was performed in PBS and allowed to react for 8 hours at room temperature and overnight at 4ºC. (**C**) Albumin-ssDNA-quencher was hybridized with biotin-ssDNA-Atto647N (complementary sequence) to assemble the Albumin-ITS (Alb-ITS).

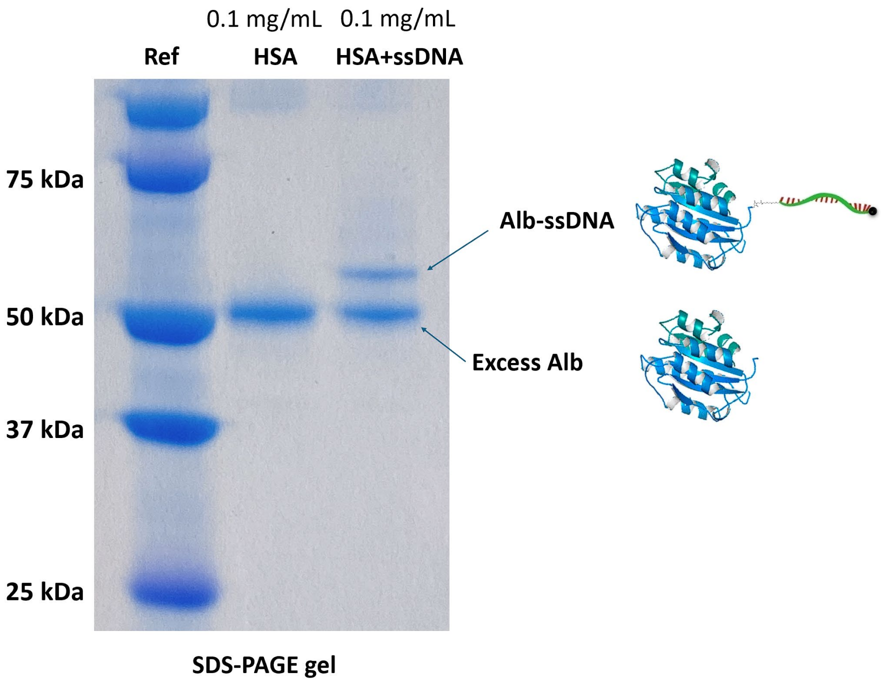


**Fig. S7. Verification of Alb-ssDNA conjugation.**

To synthesize Alb-ITS, the Alb-DNA conjugation was designed to ensure that each albumin molecule was linked to no more than one DNA strand. SDS-PAGE analysis confirmed that a fraction of human serum albumin was successfully conjugated with a single DNA strand. The excess unmodified albumin does not interfere with the preparation of the Alb-ITS surface, as it is removed during the process of Alb-ITS surface tethering through the biotin-avidin interaction.

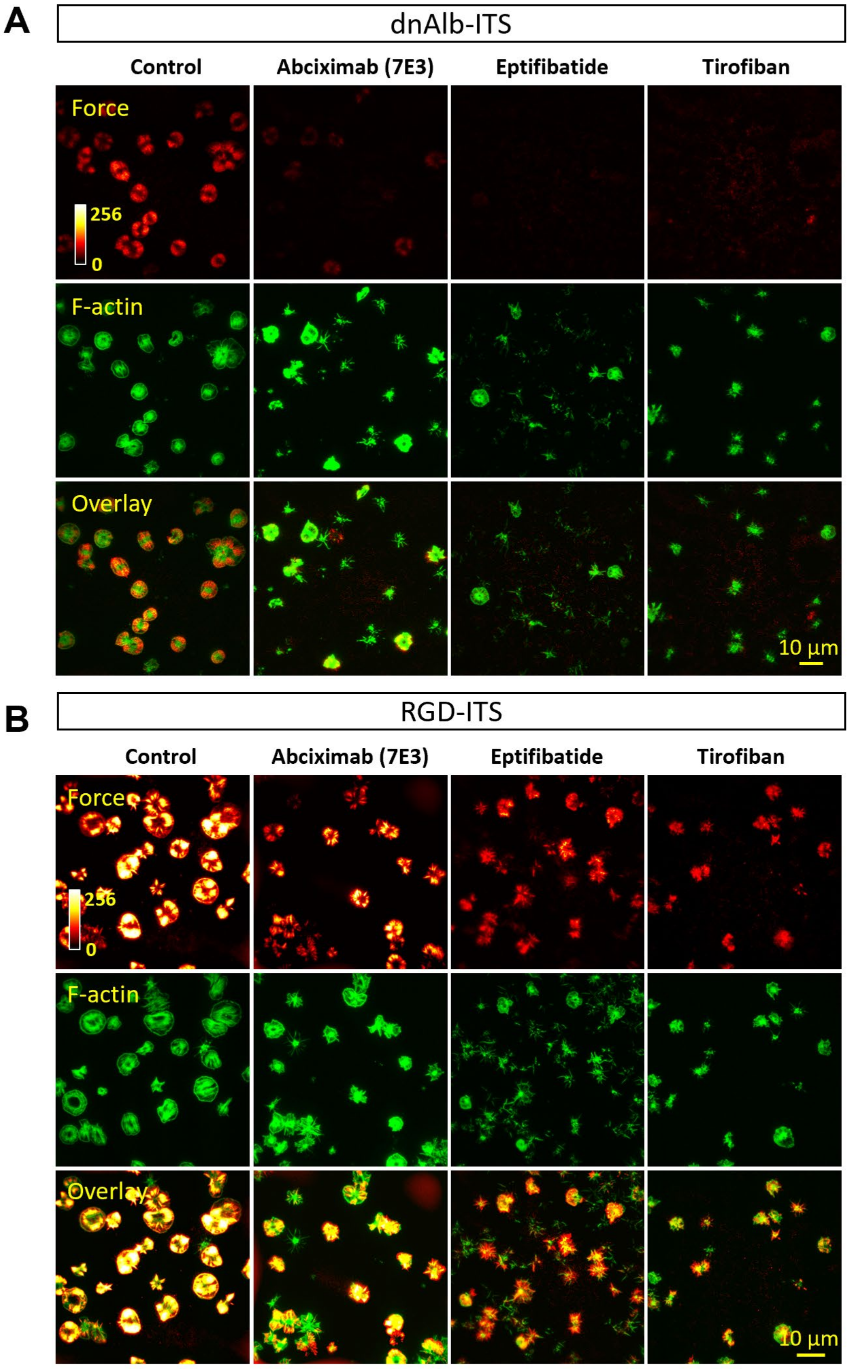


**Fig. S8. Representative images of platelet forces on dnAlb-ITS and RGD-ITS surfaces with integrin inhibition.**

(**A**) F-actin and dnAlb-transmitted platelet forces in platelets treated with three inhibitors of integrin $\alpha_{IIb}\beta_3$, 20 µg/ml Abciximab, 5 µg/ml eptifibatide and 5 µg/ml tirofiban, respectively. (**B**) F-actin and RGD-transmitted platelet forces in platelets treated with three inhibitors of integrin $\alpha_{IIb}\beta_3$.

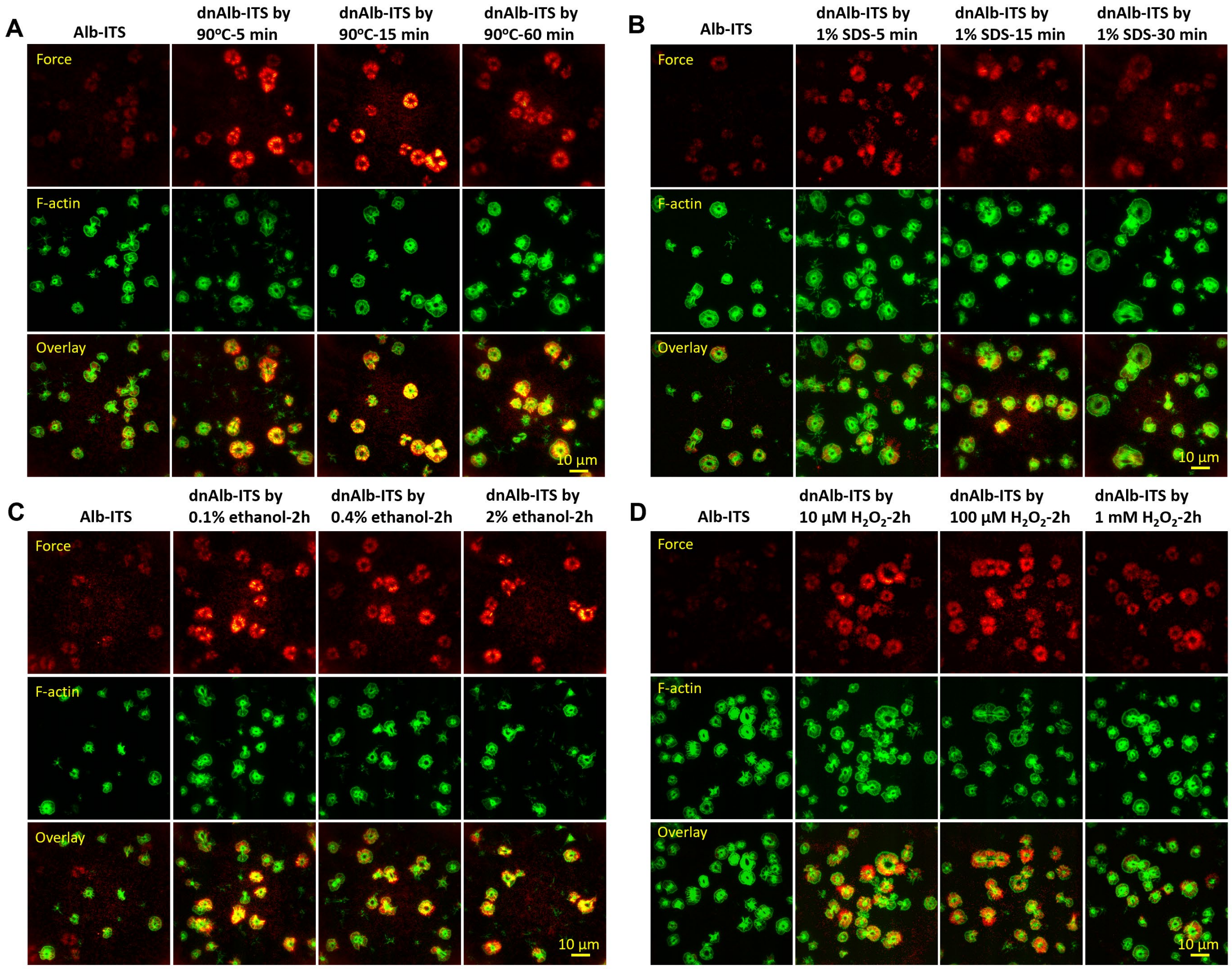


**Fig. S9. Representative platelet force images on dnAlb-ITS surfaces with the albumin denatured by various conditions**.

(**A**) dnAlb-ITS was prepared by heating Alb-ITS at 90 ºC for 5, 15 or 60 min, respectively.

(**B**) dnAlb-ITS was prepared by treating Alb-ITS with 1% SDS (sodium dodecyl sulfate) for 5, 15 or 30 min, respectively.

(**C**) dnAlb-ITS was prepared by treating Alb-ITS with 0.1%, 0.4% or 2% ethanol, respectively, for 2 h.

(**D**) dnAlb-ITS was prepared by treating Alb-ITS with 10 μM, 100 μM or 1 mM $H_2O_2$, respectively, for 2 h.

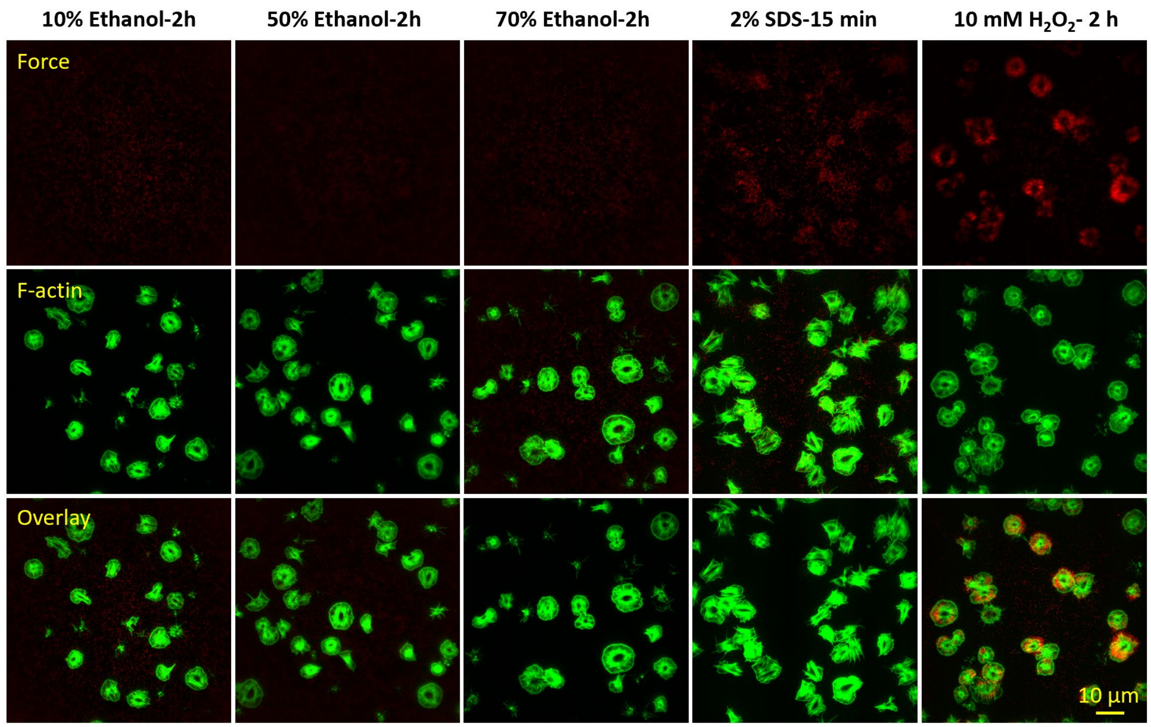


**Fig. S10. Over-denaturation of Alb-ITS led to no or low platelet force signals.**

dnAlb-ITS was prepared by treating Alb-ITS with 10% ethanol, 50% ethanol, 70% ethanol, 2% SDS or 10 mM $H_2O_2$, respectively, for 2 h. Platelet force signals become undetectable or significantly low on surfaces tethered with these dnAlb-ITS, likely due to the over-denaturation of albumin.
